# Altered composition of the *γδ* T cell pool in lymph nodes during ageing enhances tumour growth

**DOI:** 10.1101/480327

**Authors:** Hung-Chang Chen, Nils Eling, Celia Pilar Martinez-Jimenez, Louise McNeill O’Brien, John C. Marioni, Duncan T. Odom, Maike de la Roche

## Abstract

How age-associated decline of immune function leads to increased cancer incidence is poorly understood. Here, we have characterized the cellular composition of the *γδ* T cell pool in peripheral lymph nodes (pLNs) upon ageing. We found that ageing has minimal cell-intrinsic effects on function and global gene expression of *γδ* T cells, and TCR*γδ* diversity remained stable. However, ageing altered TCR*δ* chain usage and clonal structure of *γδ* T cell subsets. Importantly, IL-17-producing *γδ*17 T cells dominated the *γδ* T cell pool of aged mice - mainly due to the selective expansion of V*γ*6^+^ *γδ*17 T cells and augmented *γδ*17-polarisation of V*γ*4^+^ T cells. Expansion of the *γδ*17 T cell compartment was supported by increased Interleukin-7 expression in the T cell zone of old mice. In a Lewis lung cancer model, pro-tumourigenic V*γ*6^+^*γδ*17 T cells were exclusively activated in the tumour-draining LN and their infiltration into the tumour correlated with increased tumour size in aged mice. Thus, upon ageing, substantial compositional changes of *γδ* T cell pool in a dysregulated pLN microenvironment promote tumour growth.

## INTRODUCTION

A decline of potent T cell responses during ageing has been linked to increased susceptibility to infection and the drastic rise in cancer incidence observed in elderly mice and humans [1-3]. Three interrelated components of the immune response are affected by immunosenescence: the immune cells themselves, the supporting lymphoid organs and circulating factors that guide responses of immune cells as well as lymphoid organs [2]. In *αβ* T cells, a restricted TCR repertoire, loss of intrinsic cell functions, compromised priming as well as chronic and low-grade inflammation have been associated with impaired anti-tumour responses [1].

*γδ* T cells are unconventional T cells that combine adaptive features with rapid innate-like responses to mediate responses to infection, tissue damage, and cancer [4]. In contrast to *αβ* T cells that acquire cytokine-secreting effector functions upon activation in the periphery, murine *γδ* T cells acquire their effector potential in the thymus where they differentiate into either IFN-*γ*-producing (*γδ*1) or IL-17-producing (*γδ*17) lineages [5]. It is this pre-activated differentiation state and unique innate-like activities, that enable *γδ* T cells to rapidly infiltrate into inflammatory sites, such as tumours, in the periphery. Here they modulate the early local microenvironment and subsequent *αβ* T cell responses by secretion of pro-inflammatory cytokines [6, 7].

The anti-tumour effects of *γδ* T cells are well-established in various cancer models - mainly due to their extensive cytotoxic capacity and IFN-*γ* production [8, 9]. However, tumour-promoting roles of the IL-17-producing *γδ* T cell subsets have emerged [10, 11]. Pro-tumour mechanisms of IL-17 produced by V*γ*4^+^ and V*γ*6^+^ *γδ* T cells include the promotion of angiogenesis [12] and recruitment of immunosuppressive cells, such as myeloid-derived suppressor cells (MDSCs) [13, 14] and peritoneal macrophages [15].

Most studies on the functions of *γδ* T cells have focused on their roles in barrier tissues - mainly skin [16] and gut [17] - and in the tumour mass itself [9]. However, whether *γδ* T cells, resident in peripheral lymph node (pLN), are important for tumour-specific responses, as they have recently been shown to be for the response to inflammatory stimuli [18-21], remains unclear.

Also, currently unknown is how the *γδ* T cell pool in peripheral lymphoid tissues changes upon ageing, and how age-related alterations may affect the tumour microenvironment. For the first time, we have characterised the *γδ* T cell compartment in pLNs during ageing and investigated the functional relevance for regulating anti-tumour immune responses.

We find that, upon ageing, the *γδ* T cell pool in pLNs becomes entirely biased towards the *γδ*17 lineage, whilst *γδ*1 T cell numbers are significantly reduced. We establish that this striking *γδ*17 bias is due to a substantial accumulation of V*γ*6^+^ *γδ*17 T cells and, in part, an increased *γδ*17 polarisation of V*γ*4^+^ and V*γ*2/3/7 T cell subsets in old mice. The *γδ*17 lineage bias is supported by increased IL-7 production in the pLNs of old mice, which provides a selective niche for the expansion of *γδ*17 T cells. Interestingly, *γδ*TCR diversity is not affected, but TCR*δ* chain usage and clonal substructure is altered upon ageing. Upon tumour challenge, V*γ*6^+^ *γδ*17 T cells become activated in pLNs, migrate into the tumour and create a pro-tumour microenvironment that enhances tumour growth in the aged mice.

These results demonstrate that the *γδ* T cell pool in pLNs is essential for shaping the balance of pro- and anti-tumour immune responses. Bias towards the pro-tumourigenic *γδ*17 lineage during ageing thus may be a crucial contributor to age-related increase in tumour incidence.

## RESULTS

### γδ17 T cells constitute the majority of the γδ T cell pool in peripheral lymph nodes of old mice

To determine the effect of ageing on size and composition of the *γδ* T cell pool, we analysed inguinal and axillary lymph nodes (here termed peripheral lymph nodes, pLNs) from young (3 months old) and old (>21 months old) C57BL/6 mice. Upon ageing the proportion of *γδ* T cells amongst all CD3^+^ T cells in pLNs was increased 2-fold (Fig. 1A). The absolute number of *γδ* T cells in pLNs was slightly decreased (Fig. 1B) as a consequence of the smaller pLN size in old animals. However, when normalized to the same size, *γδ* T cells are 2 times more abundant in pLNs from aged mice (Fig. 5D, left panel). The maturation status, assessed by the characteristic lack of CD24 expression by mature *γδ* T cells, was similar between the two populations (Suppl. Fig. 1A). Thus, mature *γδ* T cells are enriched in the pLNs of old mice.

**Figure 1.**
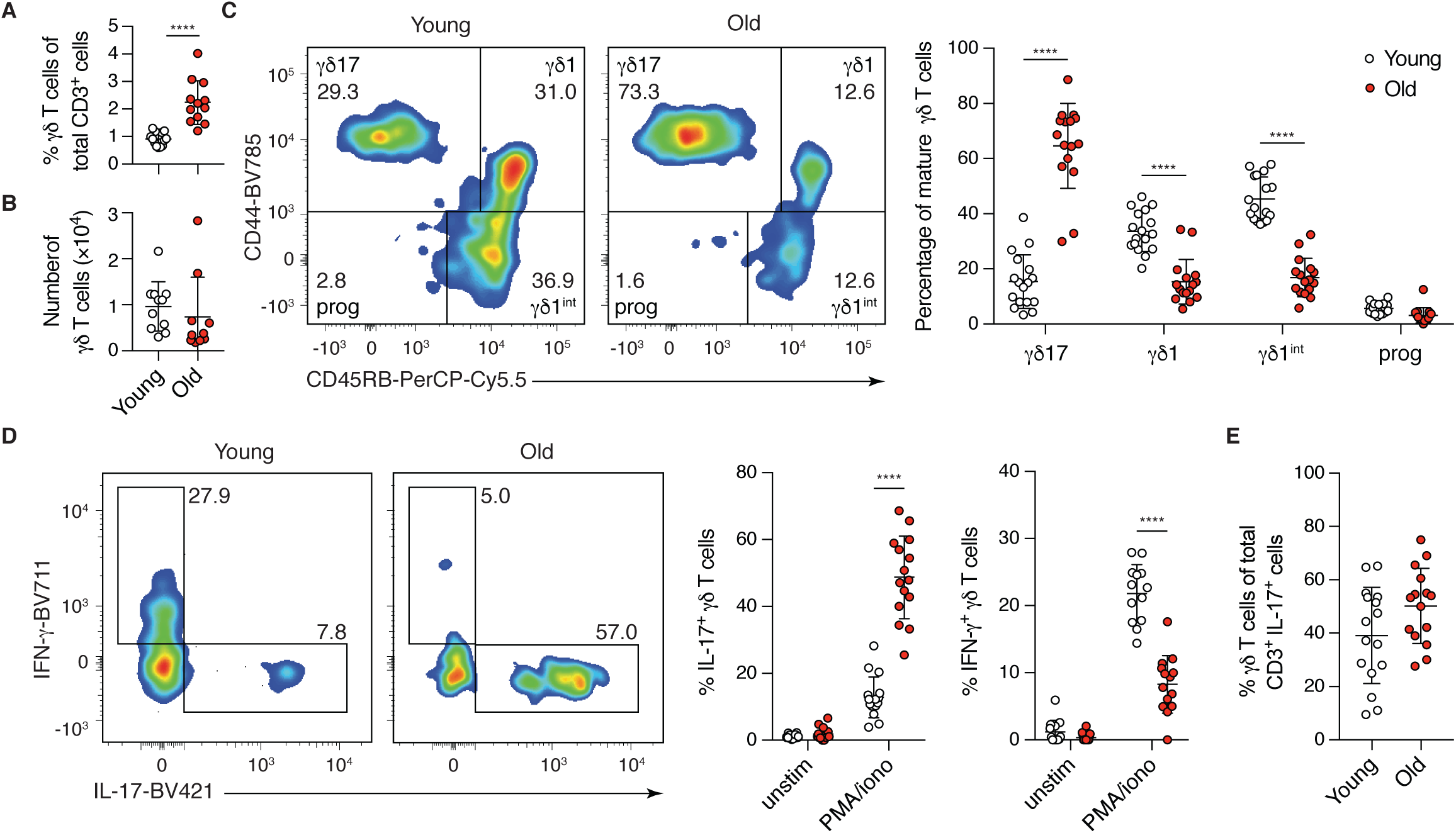
γδ T cells from peripheral lymph nodes (pLNs) of old mice are predominantly γδ17-committed. Inguinal and axillary LNs were isolated from young (3-month old, white circles) and old (>21-month old, red circles) mice. **(A)** Percentage of γδ T cells in total CD3^+^ T lymphocytes. Results are from 13 young and 12 old mice (n=5 experiments). **(B)** Absolute number of γδ T cells from pLNs in young and old mice. Results are from 11 young and 10 old mice (n=4 experiments). **(C)** CD45RB and CD44 expression of mature (CD24^-ve^) γδ T cells in pLNs of young and old mice. Left: representative FACS plots (n=11 experiments). Right: Percentage of mature γδ17-committed (CD45RB^-ve^ CD44^hi^), γδ1-committed (CD45RB^+^ CD44^+^), γδ1-intermediate (CD45RB^+^ CD44^-ve^) and progenitor (CD45RB^-ve^ CD44^-ve^) γδ T cells in pLNs of young and old mice. Results shown are from 17 young and 16 old mice (n=9 experiments). **(D)** Cell suspensions from pLNs were stimulated with PMA/Ionomycin for 4 hours and examined for their production of IL-17 and IFN-γ. Representative FACS plots are gated on CD24^-ve^ γδ T cells. **(E)** Percentage of γδ T cells in total IL-17-producing CD3^+^ T lymphocytes upon PMA/Ionomycin stimulation. Results shown (D, E) are from 16 young and 15 old mice (n=6 experiments). Statistical significances for changes in cell proportions were assessed by Mann-Whitney test (A-B), two-way ANOVA (D and F), or unpaired t test (G). \*\*\*\**p*<0.0001

In the thymus, commitment of *γδ* T cells towards *γδ*1 and *γδ*17 lineages can be distinguished by expression of CD44 and CD45RB [22]. We first confirmed that this phenotypic segregation of *γδ*1 (CD44^+^ CD45RB^+^) and *γδ*17 (CD44^hi^ CD45RB^-ve^) T cells is also observed in pLNs (Suppl. Fig. 1B). Upon stimulation with PMA/Ionomycin, CD44^hi^ CD45RB^-ve^ cells produce IL-17 but not IFN-*γ*, whereas CD44^+^ CD45RB^+^ cells produce IFN-*γ* and not IL-17. CD44^-ve^ CD45RB^+^ cells - an intermediate cell population undergoing differentiation towards the *γδ*1 lineage - produce only limited IFN-*γ* upon stimulation, and CD44^-ve^ CD45RB^-ve^ progenitor cells produce neither IL-17 nor IFN-*γ*. Consistent with previous reports [5, 10, 22-26], CD44^hi^ CD45RB^-ve^ *γδ*17 T cells were IL-7R^hi^ CCR6^+^ IL-23R^hi^ CD27^-ve^ CD62L^-ve^, while CD44^+^ CD45RB^+^ *γδ*1 T cells showed an IL-7R^lo^ CCR6^-ve^ IL-23R^lo^ CD27^hi^ CD62L^hi^ phenotype (Suppl. Fig. 1C). We next defined the contribution of *γδ*1 and *γδ*17 lineages to the *γδ* T cell pool in pLNs. We found that *γδ*1 T cells and *γδ*1 precursor (*γδ*1^int^) cells constitute greater than 80% of the *γδ* T population in pLNs of young mice whereas *γδ*17 T cells represented only 15% (Fig. 1C). Strikingly, this bias was reversed in old mice: *γδ*1 T cells are diminished and the *γδ*17 T cell population increases to 60-80% of total *γδ* T cells (Fig. 1C). pLNs from middle-aged (12 months old) animals showed an intermediate phenotype, suggesting that loss of *γδ*1 and gain of *γδ*17 T cells occurs gradually upon ageing (Suppl. Fig. 2A). We further confirmed the age-specific *γδ*1/*γδ*17 lineage redistribution using CD27 as an additional marker to separate *γδ*1 (CD27^+^) and *γδ*17 (CD27^-ve^) T cells, again observing increased proportion of *γδ*17 T cells (CD27^-ve^) in pLNs of aged mice [27] (Suppl. Fig. 2B). *γδ*17 T cells resembled highly activated T cells (CD44^hi^ CD62L^-ve^), as previously reported [24]. Interestingly, *γδ*1 T cells had a central memory-like phenotype (CD44^int^ CD62L^+^) and *γδ*1^int^ T cells showed a naïve-like phenotype (CD44^-ve^ CD62L^+^) (Suppl. Fig. 2C, D). Taken together, upon ageing the *γδ* T cell population undergoes a dramatic redistribution favouring the *γδ*17 T cell lineage.

Previous studies established that a high fat diet leading to obesity can result in a similar *γδ*17 phenotype in mice [28, 29]. The mice analysed in this study were fed a standard diet, but some old mice were obese. Importantly, both obese and lean old mice presented with a *γδ*17 bias and we observed no correlation between obesity and *γδ*17 phenotype. Moreover, our analyses of lean, middle-aged (12-month old) mice that displayed an intermediate *γδ*17 phenotype in pLNs point towards a gradual accumulation of the phenotype with age.

To determine the functional consequence of increased *γδ*17 T cells in pLNs of old mice, we assessed cytokine production upon *in vitro* stimulation with PMA/Ionomycin. While on average 10% of *γδ* T cells from young mice produced IL-17, the proportion of IL-17-producing *γδ* T cells increased to 50% in old mice. In contrast, over 20% of *γδ* T cells produced IFN-*γ* in young mice, and this decreased to below 10% of *γδ* T cells in old mice (Fig. 1D). Notably, the absolute levels of IL-17 and IFN-*γ* production by individual activated cells were similar between young and old *γδ* T cells (Suppl. Fig. 2E), indicating that, once activated, cytokine production capacity of *γδ* T cells is maintained during ageing. Despite *γδ* T cells representing only 1-2% of total T lymphocytes in pLNs, they constitute approximately half of the IL-17-producing cells upon stimulation (Fig. 1E). Taken together, we conclude that the prevalent IFN-*γ* response by *γδ* T cells in young mice becomes skewed towards an IL-17-dominated response during ageing.

### Composition of γδ T cell subsets in the pLN pool changes during ageing

Based on their TCR*γ* chain usage, *γδ* T cells can be classified into different subsets, each with distinct tissue distribution and degree of plasticity with regard to differentiation towards the *γδ*1 and *γδ*17 lineage during thymic development or in the periphery (Fig. 2A) [5, 30]. We sought to uncover the nature of the *γδ*17 bias observed in pLNs of old mice. Using the strategy described in Fig. 2B, we discriminated *γδ* T cell subsets (Heilig and Tonegawa nomenclature) [31] according to their lineage commitment. Consistent with previous reports [10, 30], V*γ*1^+^ and V*γ*4^+^ T cells were the major *γδ* T cell subsets in pLNs of young mice (Fig. 2C). By contrast, in pLNs of old mice, the V*γ*1^+^ T cell pool contracted 2-fold, and strikingly the V*γ*6^+^ T cell pool, which was barely detectable in young mice, expanded more than 10-fold. The V*γ*4^+^ T cell pool was also slightly smaller in pLNs of old mice (Fig. 2C).

**Figure 2.**
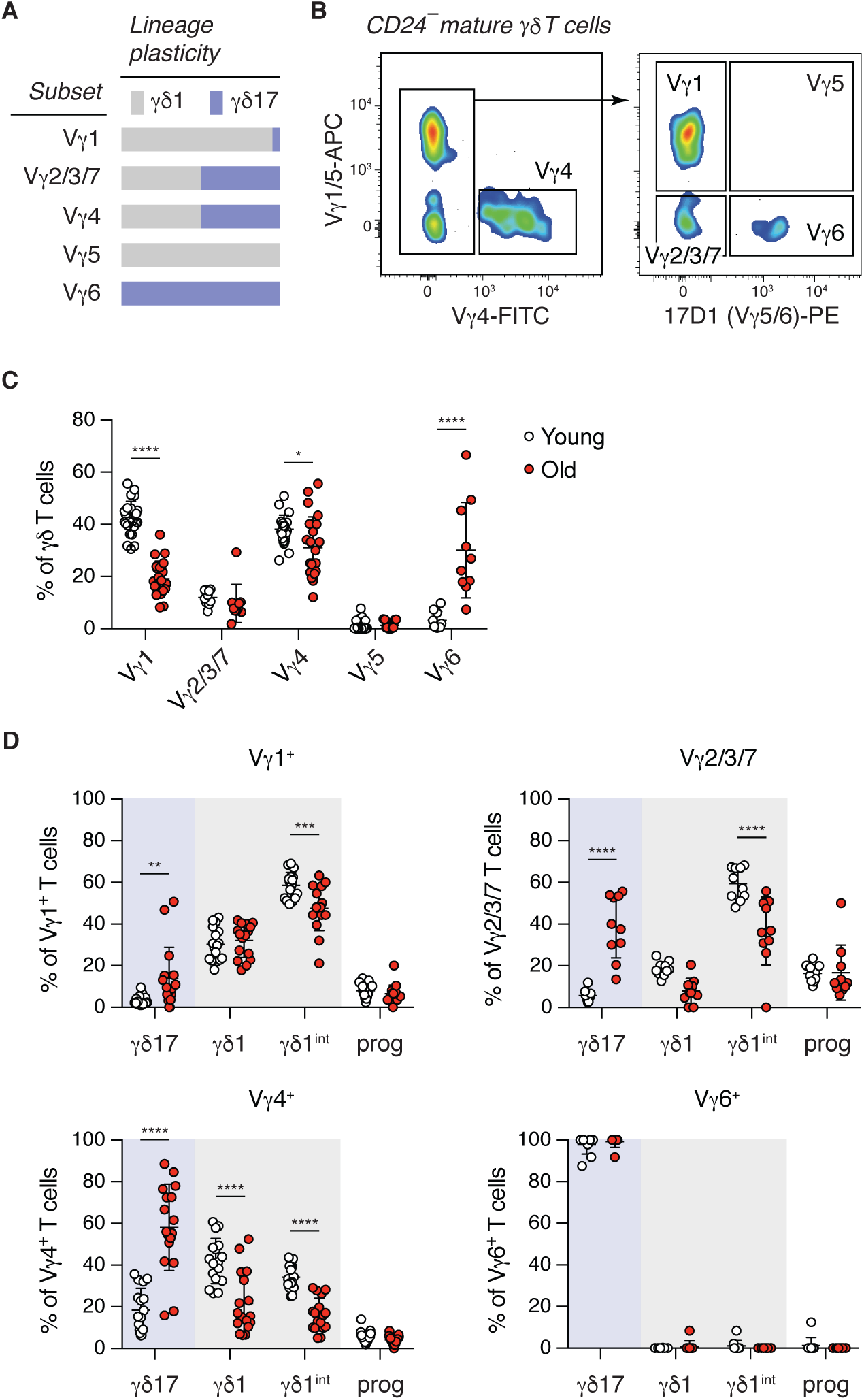
γδ17-committed Vγ4^+^ and Vγ6^+^ T cells are the main subsets in pLNs of old mice. **(A)** Published lineage plasiticity of different γδ T cell subsets according to their TCRγ chain usage. **(B)** Separation of different γδ T cell subsets according to their TCRγ chain usage by flow cytometric analysis. Expression of CD45RB, CD44 and CD27 by each γδ T cell subset was analysed (as in Fig. 1 and Suppl. Fig. 1). **(C)** Proportion of each γδ T cell subset in total γδ T cells from pLNs of young and old mice. Results shown are from 23 young and 22 old mice (n=11 experiments). **(D)** γδ1 and γδ17 lineage commitment of each γδ T cell subset in LNs of young and old mice. Results shown are from 10 pairs of young and old mice (n=6 experiments). Statistical significances for changes in cell proportions were assessed by two-way ANOVA (C and D). \**p*<0.05; \*\**p*<0.01; \*\*\**p*<0.001; \*\*\*\**p*<0.0001

V*γ*1^+^ T cells were predominantly committed to the *γδ*1 lineage in young and old mice, whereas V*γ*2/3/7 and V*γ*4^+^ T cells T cells gave rise to both *γδ*1 and *γδ*17 T cells (Fig. 2D). Although *γδ*1 T cells constitute the majority of the V*γ*2/3/7 and V*γ*4^+^ T cell pool in the pLNs of young mice, *γδ*17 cells were considerably enriched in the V*γ*2/3/7 (Fig. 2D, upper right panel) and V*γ*4^+^ (Fig. 2D, lower left panel) T cell pool in pLNs of old mice. V*γ*6^+^ T cells are invariant and exclusively committed to the *γδ*17 lineage in both young and old mice (Fig. 2D, lower right panel). Thus, enrichment of *γδ*17 lineage-committed V*γ*6^+^ T cells and changes in lineage commitment of V*γ*4^+^ and V*γ*2/3/7 T cells underpin the increase of *γδ*17 T cells in pLNs during ageing.

To determine whether the biased *γδ*17 phenotype we observe in aged mice is specific to the pLNs, we investigated the *γδ* T cell pool in the spleen, another secondary lymphoid organ. In the spleen we found a small decrease in *γδ* T cell number and the proportion of *γδ* T cells in total CD3^+^ T lymphocytes (Suppl. Fig. 3A, B), while the maturation of *γδ* T cells was unaffected by ageing (Suppl. Fig. 3C). V*γ*6^+^ T cells were also enriched, albeit to a lesser degree, thus leading to a *γδ*17 biased cell pool in the spleen of old mice (Suppl. Fig. 3D, E). The decline of the splenic V*γ*1^+^ T cell pool during ageing was less severe compared with pLNs (Suppl. Fig. 3E).

### Ageing has minimal impact on the transcriptome of γδ T cells

In order to determine the mechanism underlying the *γδ*17 bias in old pLNs, we carried out transcriptome analysis to compare purified V*γ*6^+^ *γδ*17, V*γ*4^+^ *γδ*17, V*γ*4^+^ *γδ*1, and V*γ*1^+^ *γδ*1 T cells from young and old pLNs (gating strategy provided in Suppl. Fig. 4). We confirmed purity of sorted populations by analysis of characteristic transcription factors, surface markers, cytokines, chemokines and receptors as well as effector molecules, reported to delineate respective *γδ*1 and *γδ*17 T cell subsets (Fig. 3A and Suppl. Fig. 5). Overall, when compared with *γδ*1 T cells (V*γ*1^+^ and V*γ*4^+^), *γδ*17 T cells (V*γ*4^+^ and V*γ*6^+^) showed higher expression of *Cd44* and lower expression of *Ptprc*, which are both surface markers used for the segregation of *γδ*1 and *γδ*17 T cells by FACS sorting [22]. Consistent with previous reports, V*γ*4^+^ and V*γ*6^+^*γδ*17 T cells expressed *Ccr2, Ccr6*, *Il7r* and *Il23r* at higher level and down-regulated expression of *Cd27* and *Sell* [20, 23, 24, 26, 27, 32, 33]. Master transcription factors were highly expressed in the respective lineage: *Rorc*, *Sox13*, *Maf* and *Zbtab16* in *γδ*17 T cells and *Eomes*, *Tbx21* and *Id3* in *γδ*1 T cells [18, 26, 34-41]. In homeostasis, *γδ*1 T cells expressed higher levels of *Ifng* and *γδ*17 T cells sporadically expressed *Il17a*. Interestingly, V*γ*4^+^ and V*γ*6^+^ *γδ*17 T cells expressed high levels of TCR complex component *CD3ε* [42] and of *Tcrg-C3* and *Tcrg-C1*, respectively. By contrast, cytotoxic molecules and NK receptors were highly expressed in V*γ*1^+^ and V*γ*4^+^ *γδ*1 T cells (Fig. 3A and Suppl. Fig. 5).

**Figure 3.**
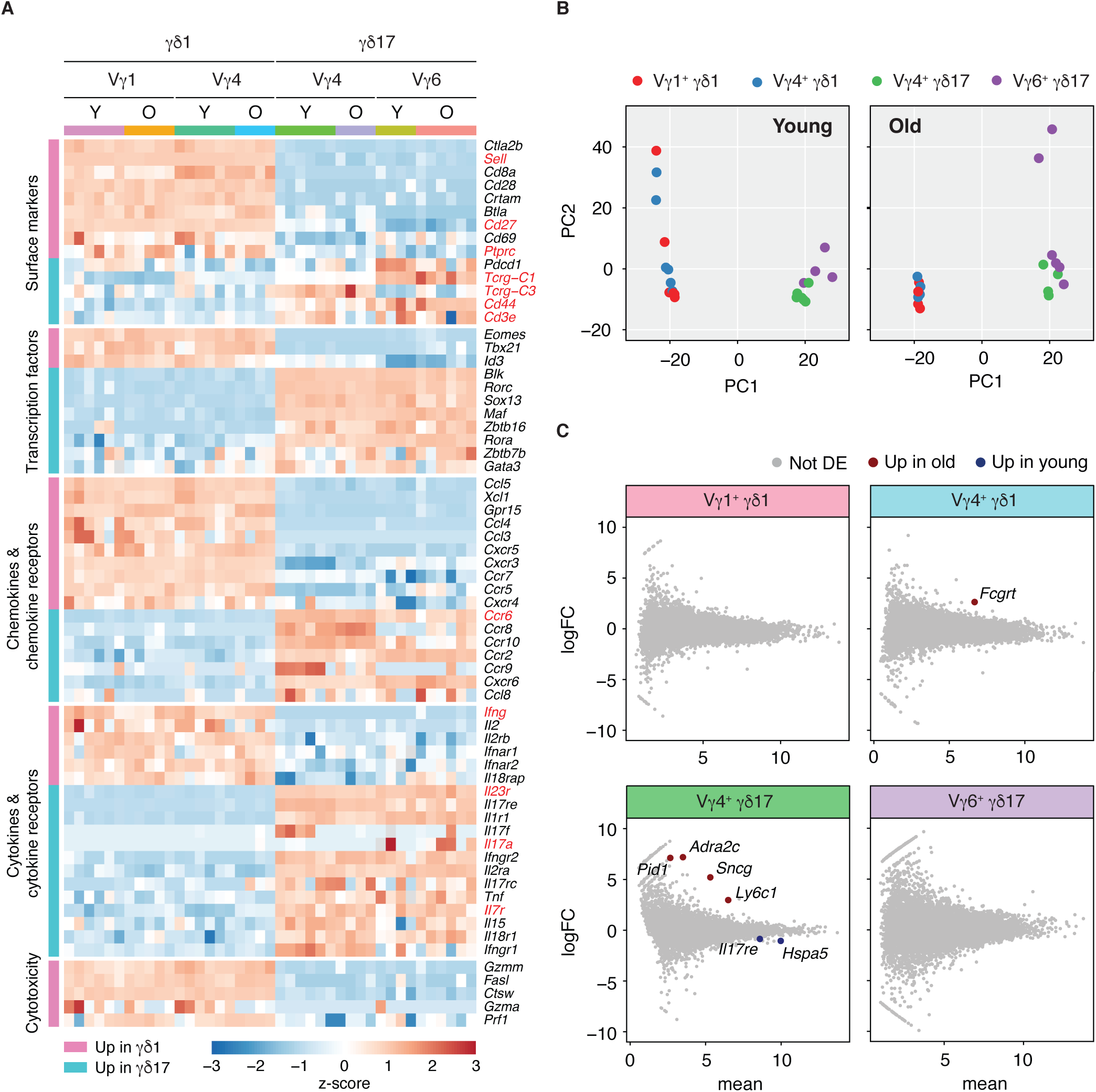
Transcriptomic analysis identifies minimal differences between γδ T cell subsets isolated from young and old mice. **(A)** Differentially expressed genes in γδ1 and γδ17 lineages were identified by RNA-Seq analysis using edgeR. Heat map of relevant genes for γδ1 and γδ17 lineage differentiation and function as well as newly identified genes. Genes shown are grouped by their functions and ranked (top to bottom) by log fold change between γδ1 and γδ17 lineages. Expression of genes marked in red were validated at the protein level by flow cytometry (IFN-γ and IL-17A was examined with or without PMA/ionomycin stimulation *in vitro*) (Fig. 1 and Suppl. Fig. 1). **(B)** Segregation of γδ1 and γδ17 T cell subsets by principle component analysis (PCA). Each dot represents one RNA-Seq library. Each library is coloured based on the cell subset. **(C)** Differential expression (DE) analysis using edgeR identifies genes up-regulated in old (red) and young animals (blue) for each subset. logFC: log fold change in expression.

Principal component analysis (PCA) revealed distinct separation between *γδ*1 and *γδ*17 lineages in PC1 but *γδ* T cells expressing different V*γ* chains were not separated in PC2 (Fig. 3B). Noteably, *γδ* T cells from young animals showed higher variance in the *γδ*1 lineage and cells from old mice showed higher variance in the *γδ*17 lineage along PC2 (Fig. 3B). Nevertheless, direct comparison of each *γδ* T cell subset from young and old mice identified only a small number of differentially expressed genes in V*γ*4^+^ *γδ*1 and *γδ*17 subsets (Fig 3C). No changes between young and old mice were detected in V*γ*6^+^ *γδ*17 and V*γ*1^+^ *γδ*1 T cells.

Since no major functional or transcriptomic changes were detected between young and old *γδ* T cell subsets we investigated whether the increase of *γδ*17 T cells in old mice could be due to (i) a change in TCR repertoire, and/or (ii) changes in the microenvironment of the pLN upon ageing.

### Ageing alters Vδ chain usage and clonal substructure but not global TCR diversity

The *αβ* TCR repertoire has been shown to decline with age [43]. We asked whether TCR diversity of *γδ* T cells from pLNs also changes upon ageing in the variant V*γ*4^+^ and V*γ*1^+^ subsets using invariant V*γ*6^+^ as a control. From the RNA-Seq data (paired-end, 125bp sequencing) of purified *γδ* T cell subsets, we reconstructed CDR3 sequences using MiXCR in the RNA-Seq mode [44, 45]. We confirmed the ability of MiXCR to reconstruct the correct V*γ* chains for each of the *γδ* T cell subsets (Suppl. Fig. 6). For further downstream analyses, we selected the *γδ* T cell subset-specific V*γ* chains. Focusing on variant V*γ*1^+^ *γδ*1, V*γ*4^+^ *γδ*1 and V*γ*4^+^ *γδ*17 cells we found surprisingly no significant difference in *γδ* TCR diversity between young and old animals, as indicated by very similar Inverse Simpson Indexes (Fig. 4A).

**Figure 4.**
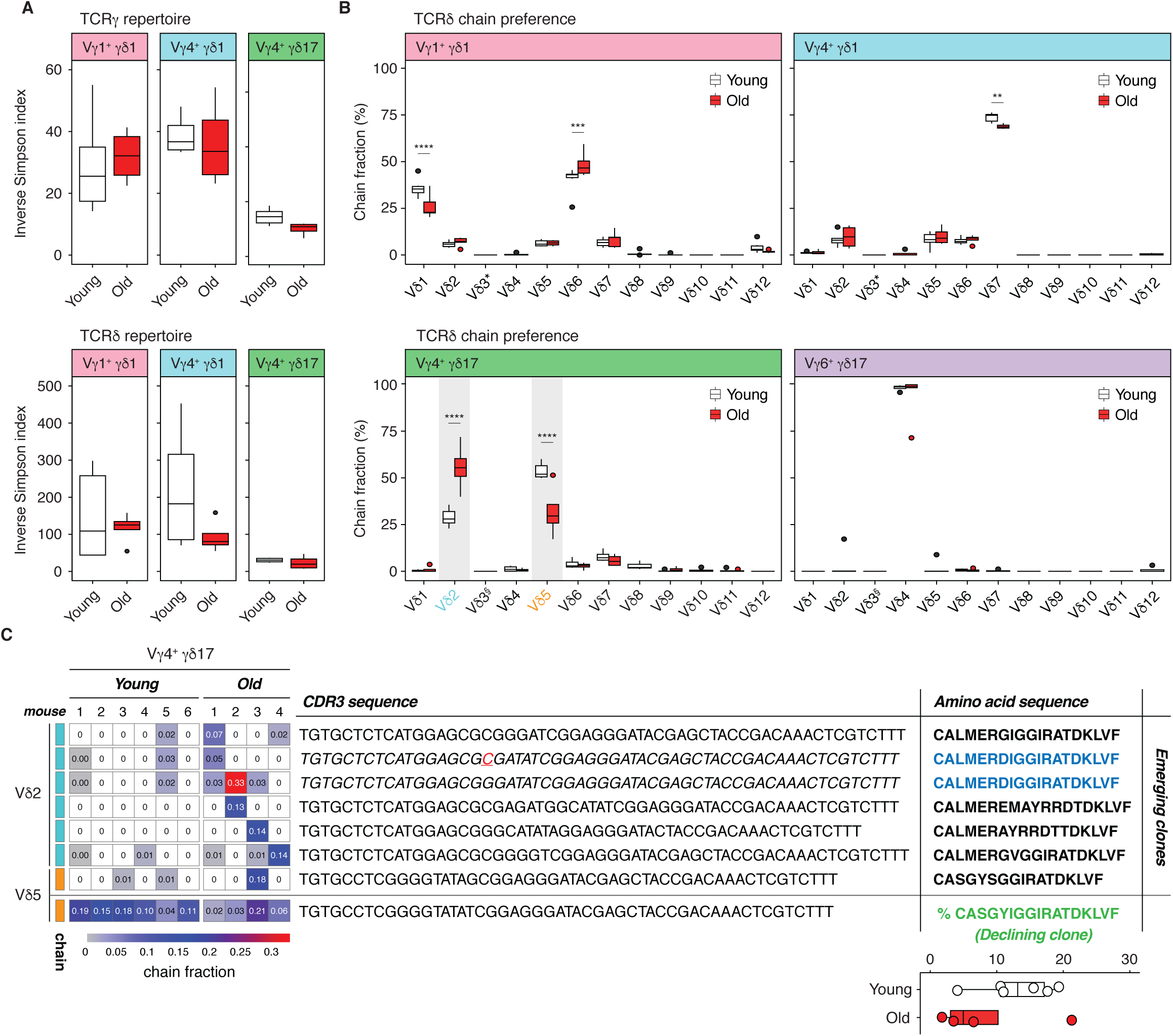
γδ TCR repertoire analysis and Vδ segment usage of γδ T cell subsets in pLNs of young and old mice. **(A)** The diversity of CRγ (top) and TCRδ (bottom) repertoires was evaluated in sorted Vγ1^+^ γδ1, Vγ4^+^ γδ1 and Vγ4^+^ γδ17 subsets of young and old mice and is epresented as inverse Simpsons index. As a control the analysis of TCRγ and TCRδ diversity in the Vγ6^+^ γδ17 subset yielded a Simpsons dex around 1 indicative of the invariant nature of this subset (data not shown). **(B)** Vδ chain usage in Vγ1^+^ γδ1, Vγ4^+^ γδ1 and Vγ4^+^ γδ17, and γ6^+^ γδ17 T cell subsets from young and old mice (^§^ Vδ3 is a pseudogene). The fraction of in-frame rearrangements of Vδ gene segments ithin the sorted populations is shown. **(C)** Emerging and declining clones defined by CDR3 nucleotide sequence in old mice. Heat map and dicated frequency show the abundance of specific clones in each young and old individual. (Inset right) Percentage of the canonical CDR3 mino-acid sequence ASGYIGGIRATDKLV in sorted Vγ4^+^ γδ17 T cells from the pLNs of young and old mice. Data shown are from 4-6 ice/condition (3 independent experiments). \*\**p*<0.01; \*\*\**p*<0.001; \*\*\*\**p*<0.0001

Next, we assessed the recombination of TCR*δ* chains and observed distinct preferences in the choice of V*δ* segments amongst the different subsets (Fig. 4B). In young mice, TCR*δ* chains utilised by V*γ*1^+^ *γδ*1 T cells contained mainly rearrangements with V*δ*6 (*∼*40%) and V*δ*1 (*∼*30%) segments, followed by V*δ*7 (*∼*10%), V*δ*5 (*∼*10%), V*δ*2 (*∼*7%), and V*δ*12 (*∼*3%), consistent with a previous study [46]. In old mice, the use of V*δ*6 increased slightly to *∼*50% and the use of V*δ*1 decreased marginally to *∼*25% compared with young mice (Fig. 4B, upper-left panel). In young mice, the majority of TCR*δ* chains from the V*γ*4^+^ *γδ*1 T cell samples contained a V*δ*7 segment (*∼*75%), followed by V*δ*5 (*∼*10%), V*δ*6 (*∼*8%), and V*δ*2 (*∼*7%) again similar to observations in a previous study [46] (Fig. 4B, upper-right panel). In old mice, the preferential use of V*δ*7 by V*γ*4^+^ *γδ*1 T cells was slightly reduced to *∼*70% (Fig. 4B, upper-right panel). Strikingly, ageing showed a profound impact on the V*δ* segment usage for TCR*δ* recombination of V*γ*4^+^ *γδ*17 T cells (Fig. 4B, lower-left panel). V*γ*4^+^ *γδ*17 T cells in young mice preferentially utilised V*δ*5 (∼50%), V*δ*2 (∼30%), and V*δ*7 segments (∼10%), for the recombination of their TCR*δ* chain. In old mice the preference among V*δ*5 and V*δ*2 segments was reversed compared with young mice: V*δ*2 (∼60%) was preferentially used followed by V*δ*5 (∼30%) (Fig. 4B, lower-left panel). As reported, invariant V*γ*6^+^ T cells from young and old mice used only the V*δ*4 segment for the assembly of their *γδ* TCR [47] (Fig. 4B, lower-right panel). The encoded CDR3 amino acid sequence (CGSDIGGSSWDTRQMFF) is the same to the V*δ*1 chain that was previously observed to be paired with V*γ*6 chain by others [42, 48, 49].

We interrogated the profound changes in TCR*δ* chain preference observed in V*γ*4^+^ *γδ*17 T cells by investigating the clonality in V*δ*2 and V*δ*5 sequences (Fig. 4C). Looking at the 10 most frequent V*δ*2 clones per individual mouse, we found 6 clones expanded (>1%) in old mice representing from >1 to up to 33% of the entire repertoire in the individual mouse. Half of the clonal expansions were private (occurring in 1 out of 4 old mice) while the other half occurred in 2-3 out of the 4 mice. When we looked at the V*δ*5 sequences we also found one expanding clone in 1 out of 4 old mice representing up to 18% of the repertoire. Furthermore, we identified two CDR3 clones from separate mice with different nucleotide sequences both giving rise to an emerging V*δ*2^+^ clone with CALMERDIGGIRATDKLVF amino acid sequence (Fig. 4C). Most interestingly, we detected the canonical ASGYIGGIRATDKLV (V*γ*4J*γ*1/V*δ*5D*δ*2J*δ*1) clone [46] in all individuals, and found that this dominant clone in young mice decreases over 50% in 3 out of 4 old mice.

Thus, although organismal ageing did not impact on global *γδ* TCR diversity, it affected V*δ* gene segment usage, led to both private and non-private clonal expansions and a collapse of the recently discovered, dominant invariant ASGYIGGIRATDKLV clone in V*γ*4^+^ *γδ*17 T cells.

### Increased IL-7 in the LN microenvironment during ageing correlates with accumulation of γδ17 T cells

To determine whether the microenvironment affects the *γδ*17 bias, we interrogated the expression of cytokines associated with activation and homeostatic maintenance of *γδ*17 T cells in primary and secondary lymphoid organs. We determined the mRNA expression levels of IL-1*β* and IL-23, which promote polarisation of *γδ*17 T cells in peripheral tissues [26, 32, 50], as well as IL-2, IL-15, and IL-7, which are involved in maintenance of *γδ* T cells [51-53], in whole pLNs (Fig. 5A) of old and young mice. Expression of IL-1*β*, IL-23, and IL-15 was not significantly different between young and old pLNs. IL-2 mRNA expression was low but slightly upregulated in old pLNs. Most strikingly, IL-7 mRNA was highly expressed in the pLNs of both young and old mice and its levels were 3-4 folds upregulated in old mice (Fig. 5A). Interestingly, IL-7 has been reported to preferentially promote the expansion of IL-17-producing CD27^-ve^ *γδ* T cell in the pLNs upon TCR stimulation [24]. As previously reported [22, 25], we found that the expression of IL-7 receptor-*α* (CD127) is over 2-fold higher in *γδ*17 compared with *γδ*1 T cells (Fig. 3A, 5B and Suppl. Fig. 1C).

**Figure 5.**
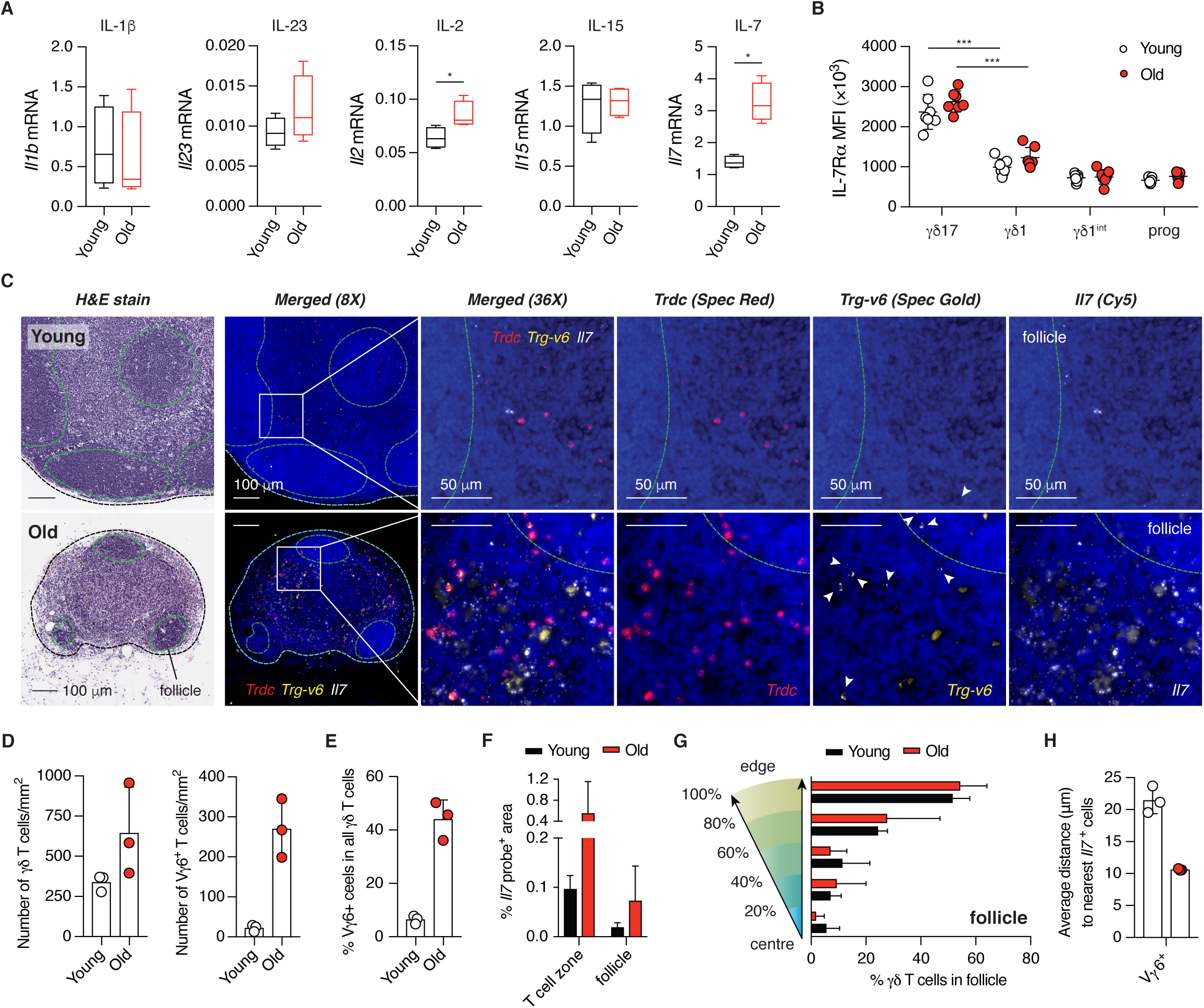
IL-7 is highly expressed in pLNs of old mice and creates a niche for V γ6^+^ γδ17 T cell expansion. **(A)** Expression of IL-1β, IL-23, IL-2, IL-15 and IL-7 mRNA in the pLNs of young (n=4) and old (n=4) mice was analysed by qRT-PCR and normalised to *Tbp* as a housekeeping gene. Similar results were obtained when using *Hprt* and *B2m* as housekeeping genes (data not shown). Results are representative of two independent experiments each with 4 young and 4 old mice. **(B)** Protein expression of IL-7Rα by γδ1 and γδ17 T cells from young and old mice. Results shown are from 7 young and 7 old mice (n=3 experiments). **(C)** Serial sections of inguinal LNs from young (top) and old (bottom) mice were stained with H&E or with specific probes targeting the constant region of TCRδ, Vγ6 TCR or IL-7 mRNA as indicated. Representative images shown are from 3 pairs of young and old mice. **(D)** Density of total γδ T cells and Vγ6^+^ T cells, **(E)** proportion of Vγ6^+^ T cells of total γδ T cells, **(F)** Expression of IL-7 mRNA in the T cell zone and the follicle of the pLN. **(G)** Localisation of γδ T cells within the follicle. **(H)** Average distance between Vγ6^+^ T cells and the nearest IL-7 producing cell. Statistical significances for changes in expression levels were assessed by Mann-Whitney test (A) or two-way ANOVA (B). \**p*<0.05; \*\*\**p*<0.001

IL-7 is constitutively secreted by stromal fibroblastic reticular cells in the T cell zone [54] and by lymphatic endothelial cells [55]. To determine whether IL-7 secreting stroma cells generate a supportive niche for V*γ*6^+^ *γδ*17 T cells during ageing, we used RNAscope^®^ to interrogate the spatial relationship between V*γ*6^+^ *γδ*17 cells and IL-7-producing cells. In young and old mice *γδ* T cells were mainly localised in the T cell zone (Fig. 5C). Despite clear involution of pLNs in aged mice, the density of *γδ* T cells in T cell zone was highly increased (Fig. 5C, D). Consistent with flow cytometric analysis (Fig. 2B), we showed that the proportion of V*γ*6^+^ *γδ*17 T cells in the *γδ* T cell pool is dramatically enriched in aged mice, suggesting the selective accumulation of this unique *γδ* T cell subset (Fig. 5E).

In pLNs, *Il-7* expression was mainly restricted to T cell zone where the expression was ∼6 fold higher compared with the follicle. Importantly, we found that the T cell zone in the old pLNs contained ∼5-fold more *Il-7* mRNA compared with young pLNs (Fig. 5F). In both old and young mice, *γδ* T cells localised to the T cell zone of the pLNs, with only a few cells found in the periphery of follicles (Fig. 5G). Strikingly, all *γδ* T cells were localised in close proximity to IL-7 mRNA expressing cells (on average <20μm) and this distance was reduced to 10μm for V*γ*6^+^ T cells in old pLN (Fig. 5H).

Taken together, we show that IL-7 production in the T cell zone of pLNs is highly increased upon ageing and correlates with the expansion of *γδ*17 T cells, especially V*γ*6^+^ T cells, resulting in a skewed peripheral *γδ* T cell pool that might favour pro-inflammatory immune responses.

### γδ17 T cell bias affects tumour development in aged mice

*γδ* T cells have important and well-established anti-tumour roles due to cytotoxic function and IFN-*γ* secretion of *γδ*1 T cells. By contrast, *γδ*17 T cells have been shown to mediate pro-tumour activities [9]. We hypothesized that LN-resident *γδ* T cells can be activated upon tumour challenge, migrate into the tumour mass and impact on the tumour microenvironment.

First, we tested whether LN-resident *γδ* T cells can infiltrate into tumours using the 3LL-A9 syngeneic Lewis lung cancer model. We blocked T cell egress from LNs by administering FTY720 to mice [19, 21]. As expected FTY720 treatment reduced the number of CD4^+^ and CD8^+^ T cells in the tumour (Suppl. Fig. 7A, B). Strikingly, also the number of *γδ* T cells was greatly reduced to below 20% upon FTY720 treatment compared with control animals (Fig. 6A). Taken together, we show for the first time that LN-resident *γδ* T cells can contribute significantly to the *γδ* T cell pool in tumours.

**Figure 6.**
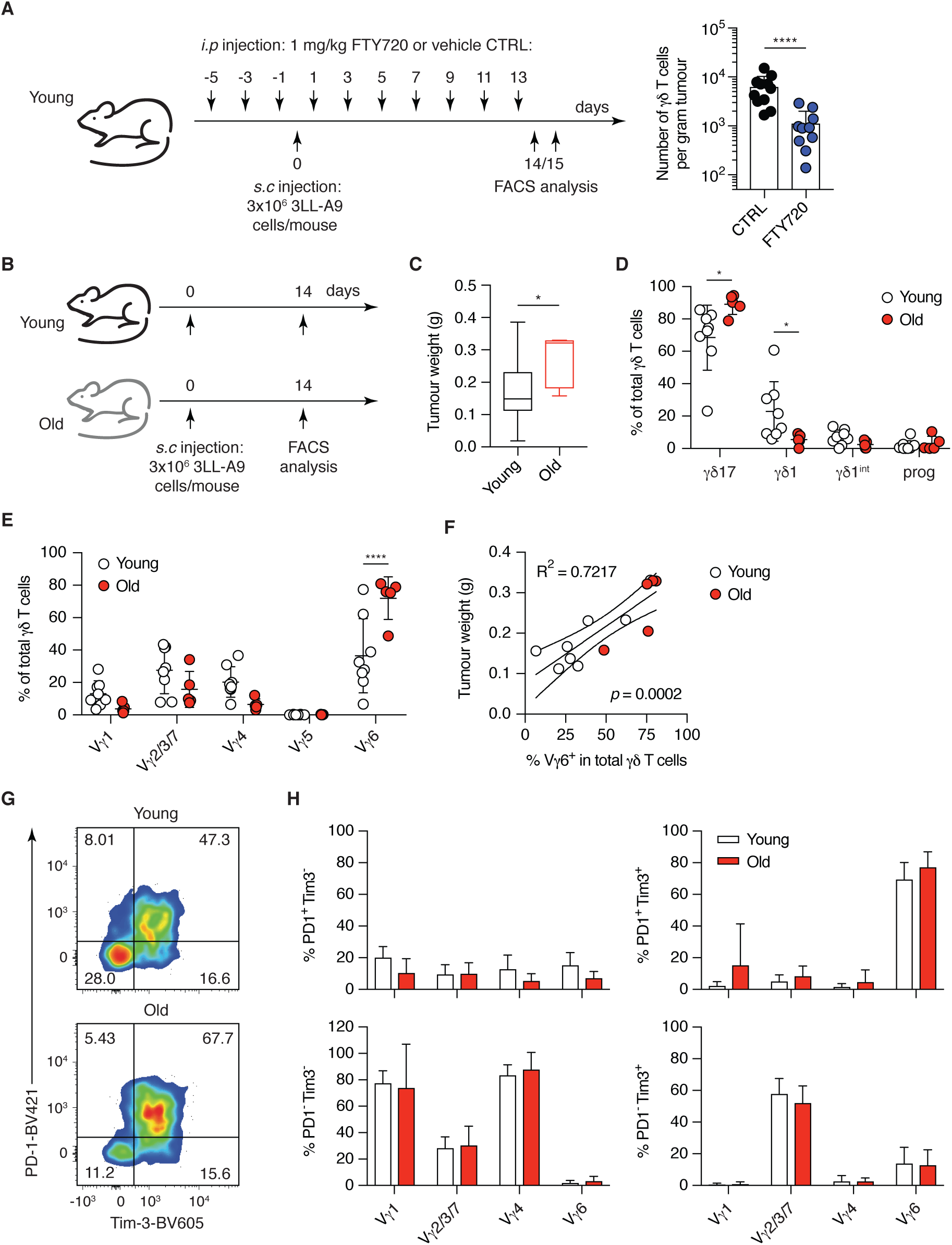
Infiltration of selectively activated V γ6^+^ γδ17 T cells from the draining LN into the tumour is associated with faster tumour progression in old mice. **(A)** Infiltration of γδ T cells from draining LN into tumour was blocked by the treatment with FTY720. Mice were injected every other day with FTY720 or vehicle control from day −5 to day 13 and 3×10^6^ 3LL-A9 Lewis lung carcinoma cells on day 0. **(B)** 3×10^6^ 3LL-A9 Lewis lung carcinoma cells were injected subcutaneously into young and old C57BL/6 mice and tumours were analysed 14 days post injection. **(C)** Tumour weights shown are from 19 young (n=5 experiments) and 5 old mice (n=2 experiments). **(D)** γδ1 and γδ17 lineage-commitment of total γδ T cells within the tumour of young and old mice. **(E)** γδ T cell subsets recovered from the tumour. **(F)** Linear regression fit between the weight of tumours and the proportion of Vγ6^+^ T cells in total tumour-infiltrating γδ T cells. **(G)** Activation and exhaustion of total tumour-infiltrating γδ T cells in young and old mice. FACS files acquired for each individual mouse were concatenated for analysis and the results are shown as representative dot plots. **(H)** Activation and exhaustion of different γδ T cell subsets in tumours from young and old mice. Results shown (D-H) are from 8 young and 5 old mice (n=2 experiments). Cell populations with the total cell number less than 10 were excluded from analysis. Statistical significances for differences were assessed by Mann-Whitney test (A and C) or two-way ANOVA (D and E). \**p*<0.05; \*\*\*\**p*<0.0001

Next, we asked whether the *γδ*17-biased *γδ* T cell pool in old mice can affect the tumour response. Strikingly, we found that 3LL-A9 tumours developed faster in old mice (Fig. 6C). *γδ* T cell infiltration into the tumour was similar in young and old mice (Suppl. Fig. 8A, B), but the balance between *γδ*1 and *γδ*17 T cells was altered: while tumours from young mice maintained a substantial proportion of anti-tumour *γδ*1 T cells, over 90% of the *γδ* T cell pool in tumours of aged mice were tumour-promoting *γδ*17 T cells (Fig. 6D). The tumour-infiltrating *γδ* T cell pool in young mice was heterogenous, containing V*γ*1^+^, V*γ*2/3/7, V*γ*4^+^, and V*γ*6^+^ T cells. In contrast, the tumour-infiltrating *γδ* T cell pool in old mice consisted mainly of V*γ*6^+^ T cells (>80%) (Fig. 6E). Importantly, the proportion of V*γ*6^+^ T cells in total tumour-infiltrating *γδ* T cell pool correlated with tumour size (Fig. 6F). Skin-resident V*γ*5^+^ T cells [56] were absent from the subcutaneous tumours of old and young mice. In the tumour, lineage bias of subsets was very different from the homeostasis observed in the pLNs. In the tumour microenvironment, progenitor and *γδ*1^int^ populations are lost, V*γ*1^+^ and surprisingly V*γ*4^+^ T cells were *γδ*1 and only V*γ*2/3/7 and V*γ*6^+^ T cells were *γδ*17-committed (Suppl. Fig. 8C).

We then asked which cells were activated and/or exhausted (by their PD1 and Tim-3 expression) in the tumour microenvironment to determine the involvement of different subsets in the anti-tumour response, (Fig. 6G, H and Suppl. Fig. 8D). Approximately 50% of *γδ* T cells in tumours of young mice and 70% of *γδ* T cells in tumours of old mice were highly activated and exhausted (PD1^+^, Tim-3^+^) (Fig. 6G). Interestingly, only *γδ*17 T cells showed high levels of activation (Suppl. Fig. 8D) while *γδ*1-committed V*γ*1^+^ and V*γ*4^+^ T cells were not activated (Fig. 6H). The majority of tumour-infiltrating V*γ*2/3/7 T cells in both young and old mice were single-positive for Tim-3^+^. Most intriguingly, only tumour-infiltrating V*γ*6^+^ T cells were highly activated/exhausted with high expression levels of PD-1 and Tim-3 (Fig. 6H).

In the tumour-draining LN, the overall lineage commitment and subset composition of *γδ* T cells were similar to the steady state in young and old mice (Suppl. Fig. 9A, B). Only the *γδ*17 bias within V*γ*2/3/7 and V*γ*4^+^ T cell subsets during ageing was not observed in the tumour-draining LN of old mice (Suppl. Fig. 9D). At the point of analysis, no activation of V*γ*1^+^, V*γ*2/3/7 and V*γ*4^+^ T cells was detected (Suppl. Fig. 9D). Importantly, only V*γ*6^+^ T cells expressed Tim-3 and PD-1 in old and young mice (Suppl. Fig. 9C). No upregulation of Tim-3 or PD-1 by V*γ*6^+^ T cells was observed in the non-tumour-draining LN in tumour bearing mice. In addition, V*γ*6^+^ T cells from pLNs of unchallenged mice did no express Tim-3 at steady state or upon short-term *in vitro* stimulation with PMA/Ionomycin (data not shown). These results suggest that V*γ*6^+^ T cells become activated in the tumour-draining LN. Interestingly, the low number of Tim-3 single positive V*γ*2/3/7 *γδ*17 T cells observed in the tumour cannot be found in the dLN, indicating a LN-independent origin of activation for this subset. These results were confirmed by the use of another syngeneic Lewis lung tumour model (Suppl. Fig. 10).

Taken together, we show that V*γ*6^+^ *γδ* T cells become selectively activated in the draining LN, migrate into the tumour and represent the majority of the tumour-resident *γδ* T cells in old mice. Furthermore, the biased *γδ*17 T cell pool in pLNs of old mice correlates with a pro-tumour microenvironment and enhanced tumour progression.

## DISCUSSION

How *γδ* T cells change upon organismal ageing has not been extensively explored. We have conducted the first comprehensive study of the murine *γδ* T cell compartment in pLNs during ageing. Remarkably, we found that - upon ageing - the *γδ*17 lineage dominates the *γδ* T cell pool at the expense of *γδ*1 T cells. The striking *γδ*17 bias with age is due, predominantly, to accumulation of V*γ*6^+^ T cells, and in part by increased *γδ*17 bias of V*γ*4^+^ and V*γ*2/3/7 T cell subsets.

The transcriptome of *γδ* T cells from young and old mice showed only minimal differences in gene expression. We found that Ly6C is upregulated in V*γ*4^+^ *γδ*17 T cells from aged mice. Previous work has determined that cross-linking of Ly6C induces LFA clustering/adhesion thereby supporting the homing of naïve and central memory CD8^+^ T cells to the lymph node [57, 58]. Whether this is a contributing factor enabling V*γ*4^+^ *γδ*17 T cells to accumulate in the pLNs requires further investigation. In addition, old *γδ* T cells secrete the same level of IFN-*γ* and IL-17 as their young counterparts. Similar results have been observed for naïve CD4^+^ T cells at transcriptional and functional level [59]. Our results thus support the notion that *γδ* T cells are intrinsically unaffected by ageing and instead the age-related *γδ*17 bias is a function of the altered environment, for instance, as we have shown, in the pLNs.

For the first time we localised *γδ* T cells, especially V*γ*6^+^ T cells, in pLNs of old mice and found that the majority of murine *γδ* T cells reside in the T cell zone. *γδ* T cells were also sporadically observed in subcapsular and medullary sinuses as previously described [60, 61] but at low level. While peripheral *γδ*1 T cells are mainly replenished through thymic output, *γδ*17 T cells are maintained persistently in the periphery [62]. Importantly, we discovered that IL-7 is highly expressed in the aged pLNs and the number of *γδ* T cells correlated with the amount of IL-7. *γδ* T cells are found in close proximity to IL-7 (on average <20μm in young and <10μm in old mice) indicating that IL-7 producing cells are creating a niche in which IL-7R*α*^hi^ V*γ*6^+^ *γδ*17 T cells can be maintained in old mice.

We next characterised the *γδ* TCR repertoire of variant subsets in young and old mice in order to elucidate changes that lead to a reduced *γδ*1 T cell pool and expansions in *γδ*17 cells. In contrast to previous work demonstrating collapse of TCR diversity in *αβ* T cells upon ageing [43], the TCR diversity does not collapse in *γδ* T cells of aged mice. TCR diversity was higher in the *γδ*1 subset compared with *γδ*17. Interestingly, V*δ* chain usage was altered especially in variable V*γ*4^+^ *γδ*17 T cells upon ageing. Analysis of the CDR3 regions of the altered V*δ* chains revealed private and semi-private clonal expansions within the V*δ*2 and V*δ*5 repertoire. Interestingly, we observed that the innate V*γ*4J*γ*1/V*δ*5D*δ*2J*δ*1 (ASGYIGGIRATDKLV) clone in the V*δ*5 repertoire [46] declined in 3 out of 4 old animals analysed. Taken together, we have shown that consequences of ageing on the *γδ* TCR repertoire are: (i) altered *δ* chain usage (ii) clonal expansions of *γδ*17 clones perhaps indicating the appearance of age-related antigens; (iii) the loss of a recently described invariant, innate clone indicating loss of a specific *γδ* T cell reactivity upon ageing.

The role of the LN-resident *γδ* T cell pool in cancer has not been explored. In tumours, *γδ* T cells are the main source of IFN-*γ* at the early stage of tumour development in young mice [6]. We asked whether the acquired *γδ*17-bias in pLNs during ageing would impact on the early tumour microenvironment. Using a Lewis lung carcinoma model and blocking egress of T cells from LNs using FTY720, we found that *γδ* T cells egressing from pLNs are constituting the majority of the *γδ* T cell pool in the tumour. Importantly, V*γ*6^+^ T cells but not any other *γδ* subset become activated in the tumour-draining LN. Due to the high numbers of these pro-tumourigenic cells in the pLNs, the tumour microenvironment becomes highly tumour-promoting and tumours progress faster in old mice. Interestingly, *γδ*17-committed V*γ*4^+^ and V*γ*2/3/7 T cells are not activated upon tumour challenge, indicating that only the invariant V*γ*6 TCR can recognise tumour-associated antigens or other signals, at least in the 3LL-A9 model.

Ageing is associated with chronic inflammation resulting from systemically increased pro-inflammatory cytokines. This predisposition to inflammatory responses can significantly affect the outcome of infection [63] and cancer immunotherapy [64]. In aged mice an increase in Th17 polarized CD4^+^ T cells [65, 66], as well as higher IL-17 secretion by liver-resident NKT cells have been described [63]. Here we discovered *γδ*17 T cells as a new critical pathogenic player during ageing.

Taken together, we have identified a novel age-dependent dysregulation of the *γδ* T cell pool that leads to enhanced tumour progression in old mice. Development of therapeutics specifically targeting *γδ*17 T cells and correcting the biased *γδ* T cell pool in the elderly might reduce the susceptibility to age-related diseases including infection and cancer.

## MATERIALS AND METHODS

### Mice

C57BL/6 mice were purchased from Charles River UK Ltd (Margate, United Kingdom) and housed under specific pathogen-free conditions at the University of Cambridge, CRUK Cambridge Institute in accordance with UK Home Office regulations. Animals were euthanized in accordance with Schedule 1 of the Animals (Scientific Procedures) Act 1986. Every mouse used was macroscopically examined externally and internally and animals with lesions or phenotypic alterations were excluded from analysis.

### Tissue processing and flow cytometry

Peripheral lymph nodes (inguinal and axillary, alone or pooled), and spleen were collected from young and old mice, respectively, mashed through a 40 μm (thymus and pLNs) or a 70 μm cell strainer (spleen) (Greiner bio-one) with the plunger of a 2 ml syringe to prepare single cell suspensions. Cells were washed with PBS once and stained with Fixable Viability Dye eFluor™780 (Thermo Fisher Scientific). Fc receptors were blocked with TruStain fcX(tm) (anti-mouse FCGR3/CD16-FCGR2B/CD32, clone 93; Biolegend) in FACS buffer containing 3% FCS (Biosera) and 0.05% Sodium Azide (Sigma-Aldrich) in Dulbecco’s phosphate buffered saline (DPBS; Gibco). Subsequently, cells were stained in FACS buffer with fluorochrome-conjugated antibodies against cell surface antigens (Suppl. Table. 1).

For the characterisation of V*γ*6^+^ T cells, the staining procedure was modified as follows. Before staining of cell surface markers, cells were stained with GL3 antibodies against TCR V*δ* followed by 17D1 hybridoma supernatant (kindly provided by Prof. Adrian Hayday, The Francis Crick Institute, London) that recognises both V*γ*5 and V*γ*6 TCR. PE-conjugated mouse anti-rat IgM monoclonal antibody (RM-7B4, eBioscience) was then used to capture cells stained positive with 17D1 hybridoma supernatant. Cells were analysed using a FACS LSR II, FORTESSA or ARIA (BD) instrument and FlowJo software (v10.4, FlowJo, LLC).

### In vitro *stimulation*

Single cell suspensions from peripheral LNs were washed twice with complete RPMI medium (RPMI-1640 (Gibco), supplemented with 10% heat-inactivated FCS (Biosera), 1 mM Sodium Pyruvate (Gibco), 10 mM HEPES (Sigma), 100 U/ml penicillin/streptomycin (Gibco) and 50 μM β-mercaptoethanol (Gibco)) and plated in 96-well plate with or without 50 ng/ml PMA (Sigma-Aldrich) and 1 μg/ml ionomycin (Sigma-Aldrich) in the presence of GolgiStop (1:1500 dilution, BD) for 4 hours. After incubation, cells were washed once with PBS and stained with Fixable Viability Dye eFluor™780 (Thermo Fisher Scientific) followed by blocking with TruStain fcX(tm) (anti-mouse FCGR3/CD16-FCGR2B/CD32, clone 93; Biolegend) and staining with fluorophore-conjugated antibodies against cell surface antigens in FACS buffer. Cells were then fixed and permeabilised using BD Cytofix/Cytoperm™Plus kit for intracellular staining with fluorochrome-conjugated antibodies against IFN-*γ* (clone XMG1.2, Biolegend) and IL-17A (clone TC11.18H10.1, Biolegend). Stained cells were run on a BD FACS LSR II cytometer and analysis was performed using FlowJo software (v10.4, FlowJo, LLC).

### Isolation of γδ1 and γδ17 T cells

Single cell suspensions were prepared from inguinal and axillary LNs collected of young and old mice. To enrich *γδ* T cells, *αβ* T cells and B cells were depleted from single cell suspensions by MACS using a biotinylated antibody against TCR*β* with anti-biotin microbeads and anti-CD19 microbeads, respectively. Enriched *γδ* T cells were then stained for FACS sorting as described above. Gating strategy used to identify V*γ*1^+^ *γδ*1, V*γ*4^+^ *γδ*1, V*γ*4^+^ *γδ*17, and V*γ*6^+^ *γδ*17 T cells is summarised in Suppl. Fig. 5. *γδ* T cell subsets were sorted with a BD FACS ARIA instrument directly into 3 μl of lysis buffer from SMART-Seq v4 Ultra Low Input RNA kit (1 μl of 10X Reaction Buffer and 2 μl of water) accordingly to the instructions of the manufacturer (Clontech). Cells were centrifuged, immediately frozen in liquid nitrogen, and stored at −80°C.

### RNA-Seq library preparation and sequencing

RNA-Seq libraries were prepared using the SMART-Seq v4 Ultra Low Input RNA kit (Clontech). Cells frozen in lysis buffers were directly complemented with cold nuclease-free water plus RNAse inhibitor (2 U/μl; Clontech) up to 9.5 μl of total volume. Volume of water was estimated calculating the number of events sorted in the BD FACS ARIA and the average drop size for the 70 μm nozzle used (∼1 nl/droplet). ERCC spike-in RNA (Ambion) (1 μl diluted at 1:300,000) and 3’ SMART-Seq CDS Primer II A (12 μM) were added to the lysis mix. cDNA was prepared following the SMART-Seq v4 Ultra Low Input RNA kit protocol (Clontech).

After cDNA preparation, RNA-Seq libraries were prepared using Illumina Nextera XT Sample Preparation Kit (Illumina, Inc., USA) and the 96 index kit (Illumina, Inc., USA). As previously described, libraries were prepared by scaling down the reactions one-fourth of the manufacturer instructions [67] and libraries were sequenced using paired-end 125bp sequencing on Illumina HiSeq4000.

### Read alignment of RNA-Seq data

Prior to read alignment, the *Mus musculus* genome (GRCm38) was concatenated with the sequence of ERCC spike-ins (available at http://tools.lifetechnologies.com/content/sfs/manuals/ERCC92.zip). Sequenced reads were aligned against this reference using *gsnap* version 2015-12-31 [68] with default settings. Gene-level transcript counts were obtained using HTSeq version 0.6.1p1 [69] with the -s option set to “no” and using the GRCm38.88 genomic annotation file concatenated with the ERCC annotation file.

### Quality control of RNA-Seq libraries

We excluded libraries with fewer than 40% of reads mapped to annotated exons or fewer than 100,000 total reads. Furthermore, we removed libraries with fewer than 10,000 genes detected with at least 1 count.

### Normalization of RNA-Seq libraries

We used the Bioconductor R package *edgeR* [70] for data normalisation. More specifically, we used the *calcNormFactors* to estimate normalisation factors and computed counts per million using the *cpm* function implemented in *edgeR*. Gene-level transcript counts are visualized as Z-score scaled, normalized counts.

### Alignment of reads to T cell receptor genes

TCR repertoire analysis was performed using the MiXCR software [44, 45]. In the first step, sequencing reads were aligned to the V, D, J and C genes of the T-cell receptor. For this, we used the *align* function with following settings: -p rna-seq -s mmu - OallowPartialAlignments=true.

### TCR assembly

The T cell receptor sequences were assembled by calling the *assemblePartial* function twice to assemble partially aligned sequences. To extend TCR alignments, the *extendAlignments* function was called. In the last step, the *assemble* function was used to fully assemble the V, D, J and C genes of the TCR.

### Exporting individual clones after TCR assembly

Individual clones were collected using the *exportClones* function while excluding out-of-frame variants (-o option) and stop codon containing variants (-t option). Clones were collected for the different chains (TRD, TRG, TRA, TRB, IGH and IGL) separately. This function returns the count, fraction as well as information on the V, D, J and C chain of the individual clones per library as defined through their CDR3 nucleotide sequence.

### Quantitative RT-PCR

Thymus, spleen and peripheral LNs (both inguinal and axillary) were collected from healthy young and old mice and homogenised with Precellys 1.4 mm Ceramic beads in 2 ml tubes (KT03961-1-003.2, Bertin Instruments) using Precellys 24 lysis and homogenisation unit (Bertin Instruments). Total RNA was extracted from homogenised samples using Ambion Purelink RNA kit (12183025, Invitrogen) according to the manufacturer’s instructions. RNA was quantitated using the NanoDrop Spectrophotometer ND-1000 and diluted in RNase-free water to 100 ng/μl for analysis. Quantitative RT-PCR was carried out using the Superscript III Platinum One-Step qRT-PCR Kit (Life Technologies) and TaqMan™Gene Expression Assays (Fam) (Life Technologies) to quantify the expression of following genes: *Tbp* (assay ID: Mm00446971_m1), *Hprt* (Mm03024075), *B2m* (Mm00437762_m1), *Btn1a1* (Mm00516333_m1), *Btnl1* (Mm01281669_m1), *Btnl2* (Mm01281666_m1), *Btnl4* (Mm03413106_g1), *Btnl6* (Mm01617956_mH), *Btnl9* (Mm00555612_m1), *Skint1* (Mm01720691_m1), *Il1b* (Mm00434228_m1), *Il2* (Mm00434256_m1), *Il7* (Mm01295803_m1), *Il15* (Mm00434210_m1), *Il17a* (Mm00439618_m1 IL17a), and *Il23a* (Mm00518984_m1). qRT-PCR was performed using a QuantStudio 6 Flex Real-Time PCR System (Thermo Fisher). For reverse transcription the thermal cycler was set at 50°C for 15 minutes followed by a 2 minutes incubation at 95°C, after which 50 PCR cycles of 15 seconds at 95°C followed by 1 minute at 60°C were run. All samples were run in triplicates and similar results were obtained for all housekeeping genes used (*Tbp, Hprt, and B2m*).

### RNAscope^®^

Inguinal LNs were isolated from young and old mice, respectively, fixed in 10% NBF (Pioneer Research chemicals Ltd) for 24h, transferred into 70% ethanol for 24h, and embedded into paraffin blocks. Paraffin sections were cut at 3mm onto Superfrost plus slides and baked for 1 hour at 60°C. Probes and Kits (RNAscope^®^ LS Multiplex Reagent Kit, Cat# 322800 and RNAscope^®^ LS 4-Plex Ancillary Kit Multiplex Reagent Kit Cat# 322830) were obtained from Advanced Cell Diagnostics. TSA Plus Fluorescein System for 50-150 Slides (Cat# NEL741001KT), TSA Plus Cyanine 3 System for 50-150 Slides (Cat# NEL744001KT), TSA Plus Cyanine 5 System for 50-150 Slides (Cat# NEL745001KT), and Opal 620 Reagent Pack (Cat# FP1495001KT) were from Perkin Elmer. Probes (automated Assay for Leica Systems) and reference sequences were as follows: RNAscope^®^ LS 2.5 Probe-Mm-Il7, GenBank: NM_008371.4 (2-1221), RNAscope^®^ 2.5 LS Probe-Mm-Tcrg-V6, GenBank: NG_007033.1: (2-475), and RNAscope^®^ LS 2.5 Probe-Mm-Trdc, GenBank: gi|372099096 (9-1098). Fluorescein staining was used as a dump channel for the exclusion of cells with unspecific background staining. Different combinations of fluorochromes were used for each probe to avoid bias in staining. Slides were scanned with Axio Scan (Zeiss) and images were analysed using Halo software (Indica Labs).

### In vivo *tumour model*

3LL-A9 cells were grown in DMEM (Gibco) supplemented with 10% FCS and tested negative for mycoplasma (MycoProbe^®^ Mycoplasma Detection Kit, R&D systems) and mouse pathogens (M-LEVEL 1 analysis, Surrey Diagnostics). For injection, the right flank of young and old mice was shaved and 3 *×* 10^6^ 3LL-A9 Lewis lung cancer cells were injected subcutaneously. Mice were sacrificed on day 14 post inoculation and the tumour and tumour-draining LN were harvested for characterisation of *γδ* T cells by flow cytometry. FTY720 (Sigma) was reconstituted in ethanol and diluted in 2% *β*-cyclodextrin (Sigma) for injections. Mice were injected *i.p* every other day with FTY720 at 1 mg/kg or with vehicle control 3 times before *s.c* injection of 3×10^6^ 3LL-A9 cells at right flank. FTY720 treatment was continued on day 1 post tumour cell injection for further 7 times before tissue collection at day 14/15.

Tumour tissue was weighed and minced by surgical curve scissors and then mashed through a 70 μm cell strainer (Greiner Bio-one) with the plunger of 2 ml syringe. Flow-through was passed again through a 40 μm cell strainer (Greiner bio-one) to prepare single-cell suspension. Immune cells were subsequently enriched from cell suspension by gradient centrifugation using Optiprep™density gradient medium (Sigma-Aldrich). Briefly, cells were resuspended in 10 ml 33.3% Optiprep™(diluted with PEB containing PBS with 0.5% BSA and 5 mM EDTA) and 5 ml PEB were layered gently on top of cell suspension without disturbing the interface of two layers. Cells were then centrifuged at 500xg for 20 minutes at 4°C without brake in the end of centrifugation. Immune cells at the interface between two layers were collected and washed twice with PBS before flow cytometric analysis.

### Statistical analysis

Statistical analysis was performed using Prism 7 software (GraphPad Inc.). Each data set was firstly analysed by D’Agostino & Pearson normality test for Gaussian distribution. Unpaired t test was used for the comparisons between two data sets (young *vs* old) both with normal distribution. Comparisons between two groups (young *vs* old) failed to pass normality test were performed using Mann-Whitney test. Two-way ANOVA with Sidak Multiplicity Correction test was used to compare multiple variables, such as *γδ* T cell lineages and subsets, between two different groups (young *vs* old). Descriptive statistics are expressed as mean *±* SD (standard deviation) in all figures. All statistical analyses were performed as two-tailed tests, and the level of statistical significance in differences was indicated by *p* values in all figures (\**p*<0.05; \*\**p*<0.01; \*\*\**p*<0.001; \*\*\*\**p*<0.0001).

### Statistical analysis for RNA-Seq data

Principal component analysis of normalized, log_10_-transformed counts was performed using the *prcomp* function in R. Differential expression analysis was performed using the Bioconductor package *edgeR* [70]. A quasi-likelihood negative binomial generalized log-linear model was fitted to the count data after removing lowly expressed genes (averaged expression <10 counts). The *glmQLFTest* function was used to perform genewise statistical testing incorporating the age of the animals as contrasts. Gene-level differential expression tests with a false discovery rate smaller than 10% were considered as statistically significant. To profile clonal diversity within each library, we calculated the inverse Simpsons index of the clone count as implemented in the R package *tcR* [71]. To allow the comparisons between libraries, we subsampled the clones to similar numbers within each T cell subset.

## ACKNOWLEDGEMENTS

Special thanks and gratitude go to Julia Jones and Cara Brodie from the Histopathology/ISH Core at the CRUK Cambridge Institute for their support in conducting RNAscope^®^ experiments and analysis; Angela Mowbray and Matthew Clayton from BRU for expert animal care; and the Flow cytometry core for cell sorting. We thank Prof. Adrian Hayday and Dr. Anett Jandke at King’s College London for supplying 17D1 hybridoma supernatant and a FACS staining protocol. We also thank Dr. Matthias Eberl at Cardiff University for constructive comments on the manuscript.

This work has been funded by the Cancer Research UK (H-CC, LMOB, DTO, JCM, and MdlR), Wellcome Trust Sir Henry Dale Fellowship (WT107609, MdlR), Janet Thornton Fellowship (WT098051, CPJM), European Research Council (DTO), EMBO Young Investigators Programme (DTO), EMBL (NE and JCM), Wellcome Trust Sanger Institute (CPMJ, JCM, and DTO) and NIHR pump priming grant (H-CC).

## AUTHOR CONTRIBUTION

HCC and MdlR designed the experiments; HCC, CPMJ and LMOB performed experiments and experimental analyses; NE performed computational analyses; LMOB provided technical assistance and support; DTO provided the ageing colony. HCC and MdlR wrote the manuscript. All authors commented on and approved the manuscript.

## DATA AVAILABILITY

The RNA-Seq data will be made available on ArrayExpress under the accession number E-MTAB-7178. Analysis scripts for the RNA-Seq and TCR analysis can be found at: https://github.com/MarioniLab/GammaDeltaTcells2018

## Supplementary Table and Figures

**Suppl. Fig. 1.**
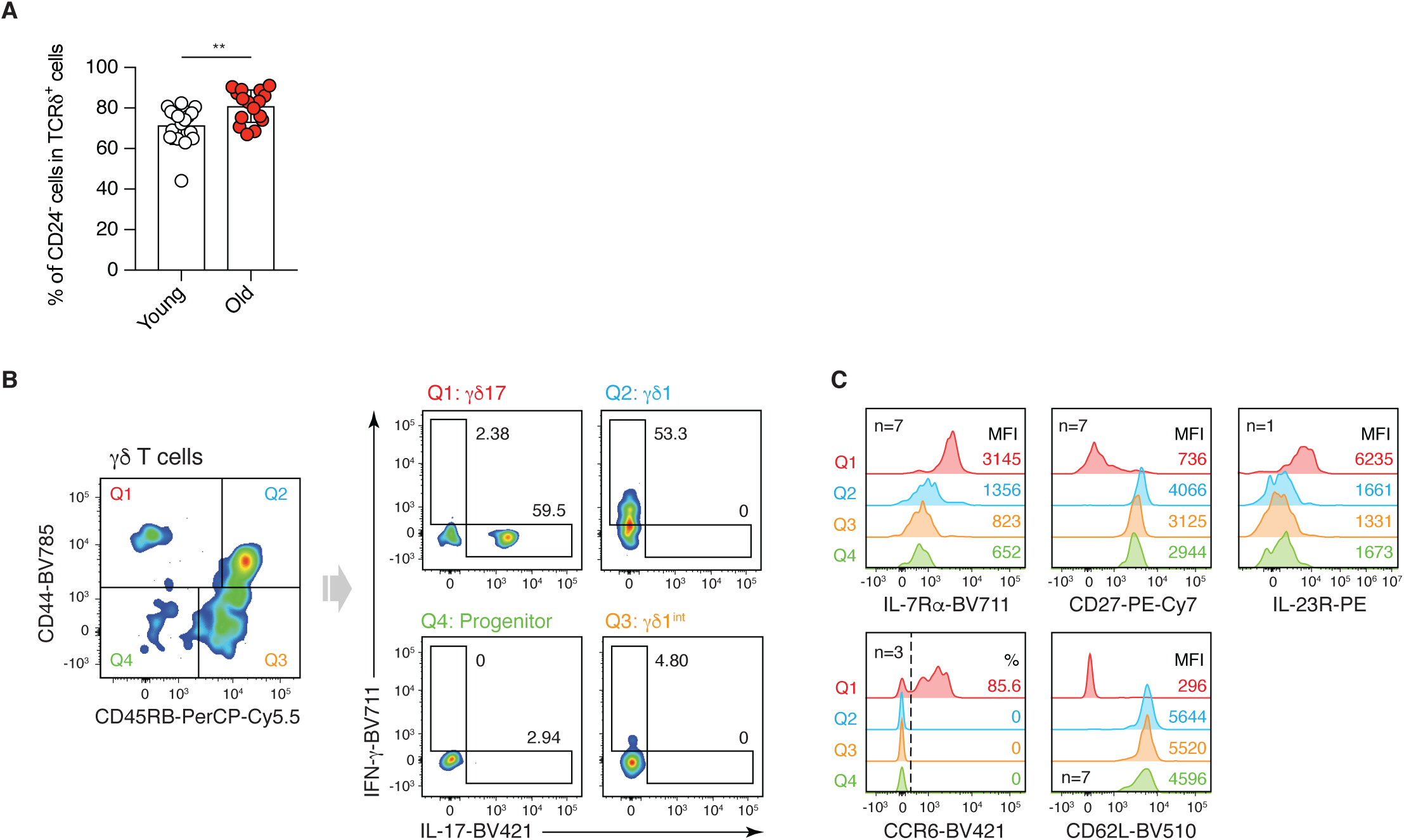
CD44 and CD45RB expression by *γδ* T cells in peripheral lymph nodes (pLNs) identifies IL-17-producing (*γδ*17) and IFN-*γ*-producing (*γδ*1) lineages. **(A)** Maturation status of γδ T cells in pLNs of young and old mice according to expression level of CD24. Results shown are from 8 independent experiments with 17 young and 16 old mice. **(B)** γδ T cells were harvested from pLNs of young and old mice, and stimulated with 50 ng/ml PMA and 1 μg/ml ionomycin in the presence of GolgiSTOP. After 4 hours, cells were stained with antibodies against cell surface markers and intracellular IL-17 and IFN-γ. FACS plots are representative of 6 independent experiments with 16 young mice. γδ T cells isolated from old mice showed similar results (not shown). **(C)** Expression of characteristic γδ1 and γδ17 lineage makers by populations separated via CD44 and CD45RB expression. FACS plots shown are representative for the number of mice indicated.

**Suppl. Fig. 2.**
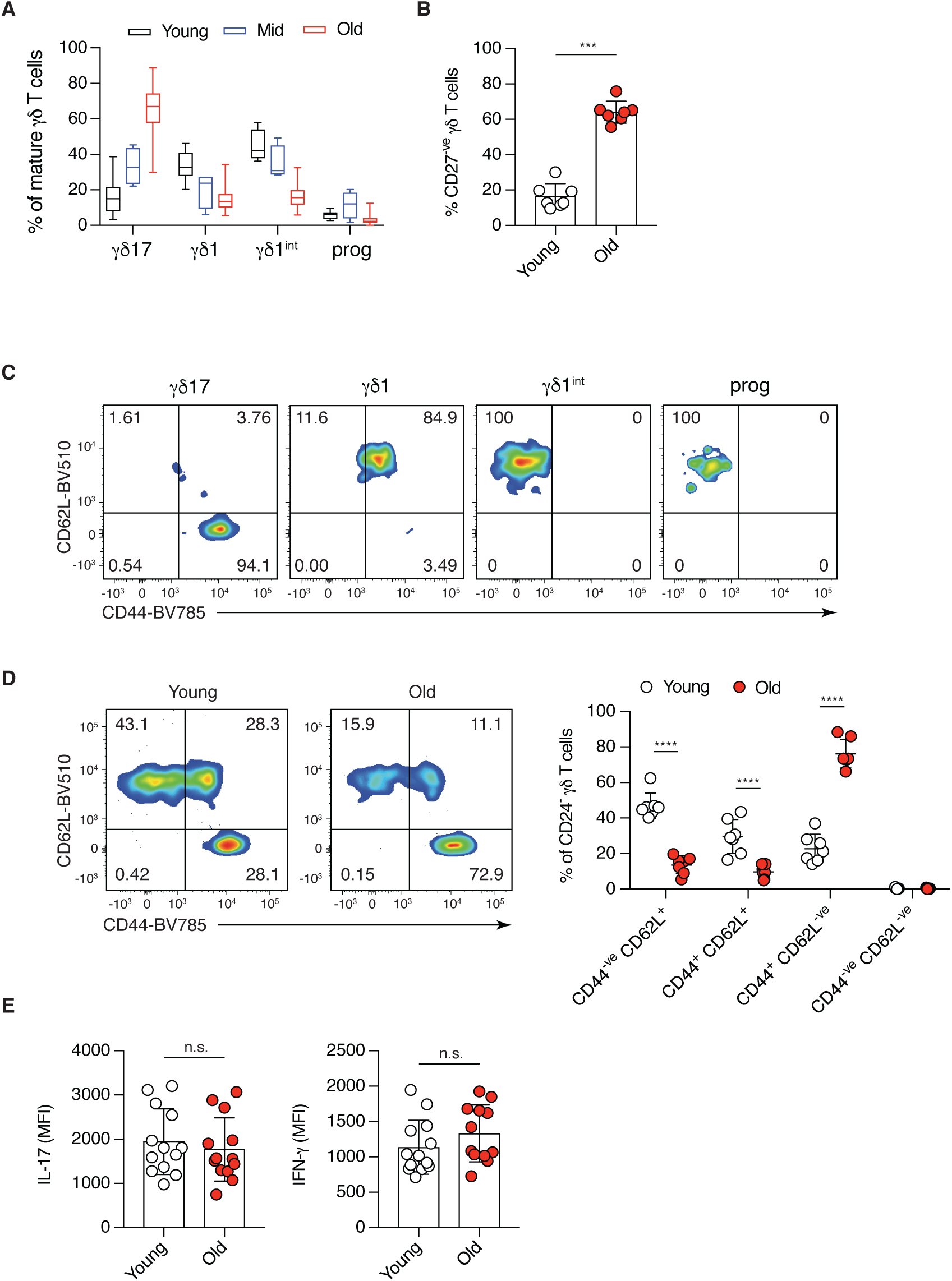
Lineage polarisation, phenotype and function of *γδ* T cells in the pLNs of young and old mice. **(A)** CD45RB and CD44 expression segregates mature (CD24^-ve^) γδ T cells in LNs of young (3 months, n=17), mid-age (12 months, n=4) and old (> 21 months, n=16) mice into γδ17-committed (CD45RB^-ve^ CD44^+^), γδ1-committed (CD45RB^+^ CD44^+^), γδ1-intermediate (CD45RB^+^ CD44^-ve^) and progenitor (CD45RB^-ve^ CD44^-ve^) γδ T cell subsets. **(B)** Percentage of γδ17 T cells, characterized by lack of CD27 expression, from total γδ T cells in old and young pLNs. Results shown are from 3 independent experiments with 7 young and 7 old mice. **(C)** CD44 and CD62L expression profile of γδ17-committed (CD45RB^-ve^ CD44+), γδ1-committed (CD45RB^+^ CD44^+^), γδ1-intermediate (CD45RB^+^ CD44^-ve^) and progenitor (CD45RB^-ve^ CD44^-ve^) γδ T cells from pLNs of young mice. **(D)** Representative FACS plots and analysis of the memory/activation status of γδ T cells in the pLNs of young and old mice. Results shown are collected from 3 independent experiments using 7 young and 7 old mice. **(E)** IL-17 and IFN-γ production by γδ T cells from old and young mice. γδ T cells were harvested from pLNs and stimulated with PMA and ionomycin for 4 hours in the presence of GolgiSTOP. Results shown are from 6 independent experiments with 16 young and 15 old mice. Statistical significance for changes were assessed using Mann-Whitney test (A and E) or two-way ANOVA (D). \*\*\**p*<0.001; \*\*\*\**p* <0.0001

**Suppl. Fig. 3.**
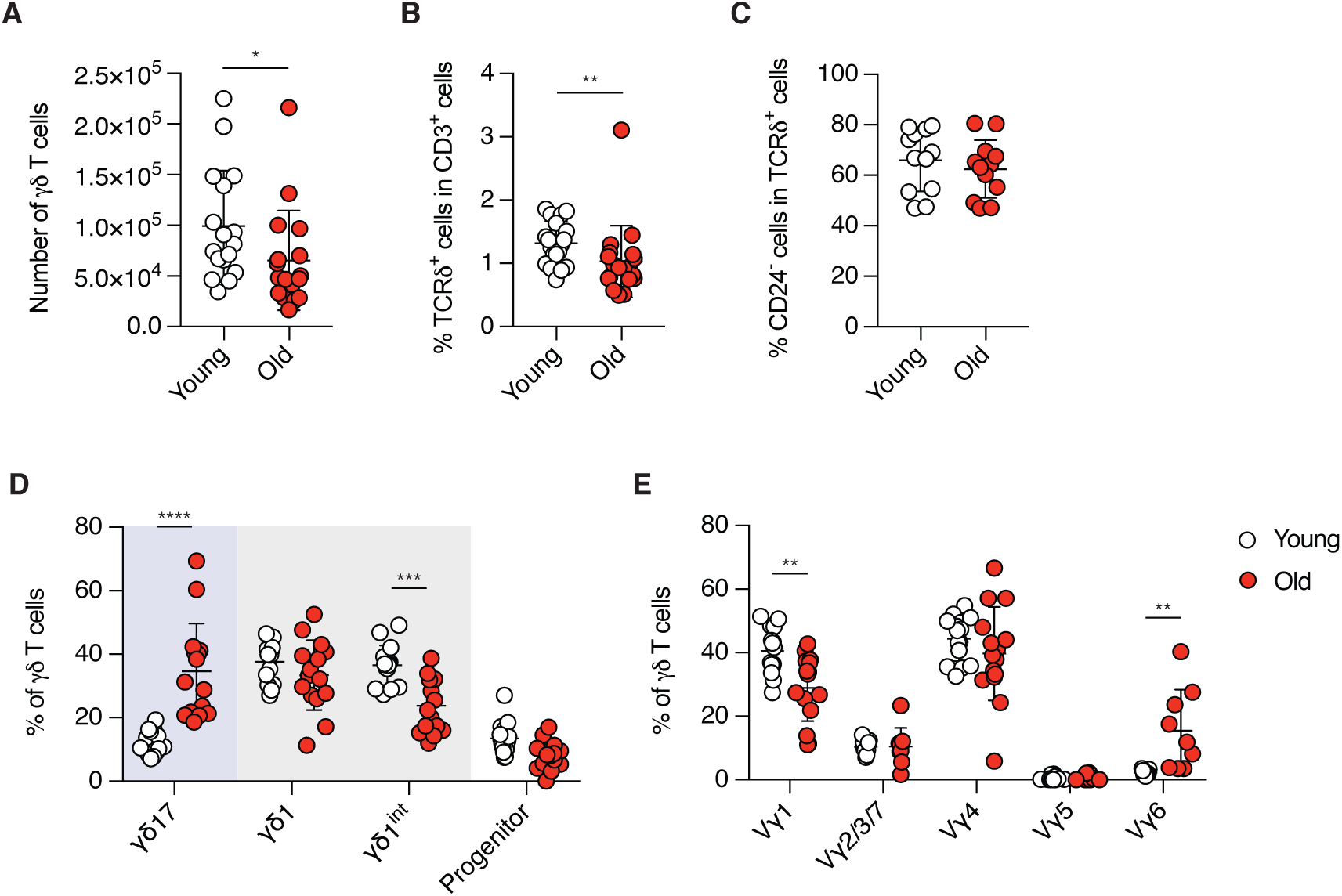
Characterisation of the splenic *γδ* T cell pool in young and old mice. **(A)** Absolute numbers of γδ T cells in the spleen of young and old mice. **(B)** Proportion of γδT cells in total CD3^+^ T lymphocytes in the spleen of young and old mice. **(C)** Maturation status, as determined by CD24 expression, of γδ T cells in the spleen of young and old mice. **(D)** γδ1 and γδ17 lineage commitment and **(E)** proportion of γδ T cell subset of mature γδ T cells in the spleen of young and old mice. Results shown are collected from 7 independent experiments with 17 young and 17 old mice. Statistical significance for difference was assessed by Mann-Whitney test (A-C) or two-way ANOVA (D and E). \**p*<0.05; \*\**p*<0.01; \*\*\**p*<0.001; \*\*\*\**p*<0.0001

**Suppl. Fig. 4.**
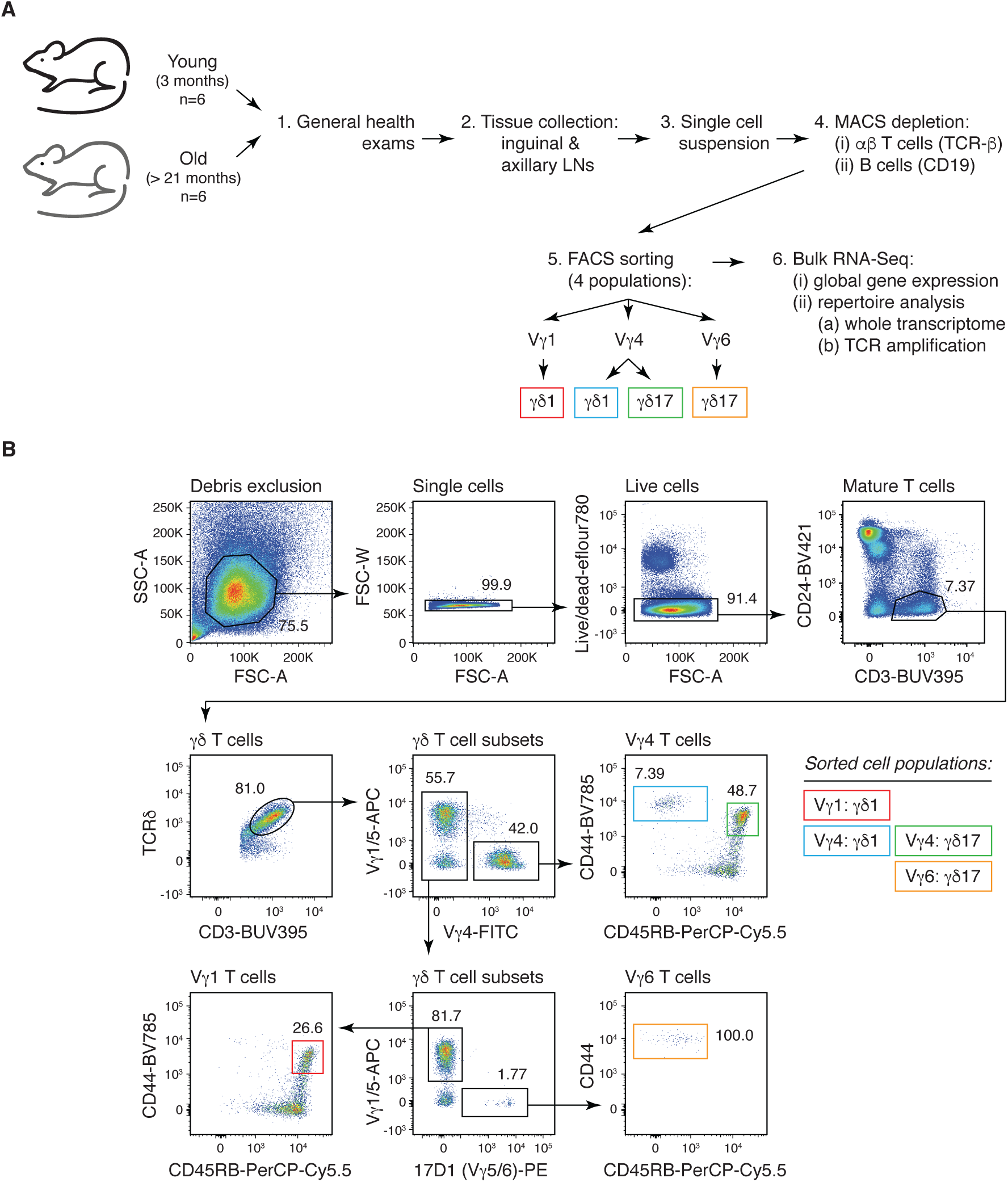
Isolation of *γδ*1 and *γδ*17 T cells from different *γδ* T cell subsets from pLNs of young and old mice. **(A)** Four populations of γδ T cells were isolated from the pLNs of young and old mice for RNA-Seq. **(B)** FACS gating strategy: Lymphocytes were gated by forward scatter (FSC-A) and side scatter (SSC-A). Cell doublets were excluded according to area and width of the forward scatter (FSC-A/FSC-W). Dead cells were removed using viability dye. From mature T lymphocytes (CD3^+^ CD24^-ve^), γδ T cells were determined by TCRδ expression. γδ T cells were then further segregated into 4 cell subsets according to their expression of different TCRγ chains. Vγ1^+^, Vγ4^+^ and Vγ6^+^ T cells were separated by the staining profile of cells with antibodies against Vγ1^+^, Vγ4^+^ and Vγ5^+^ TCR and 17D1 hybridoma supernatant against Vγ5/Vγ6 TCR. γδ1 (CD44^+^ CD45RB^+^) and γδ17 (CD44^hi^ CD45RB^-ve^) T cells were characterised within Vγ1^+^, Vγ4^+^ and Vγ6^+^ T cells. γδ1 T cells were isolated from Vγ1^+^ (red) and Vγ4^+^ (blue) T cell subsets and γδ17 T cells were isolated from Vγ4^+^ (green) and Vγ6^+^ (orange) T cell subsets.

**Suppl. Fig. 5.**
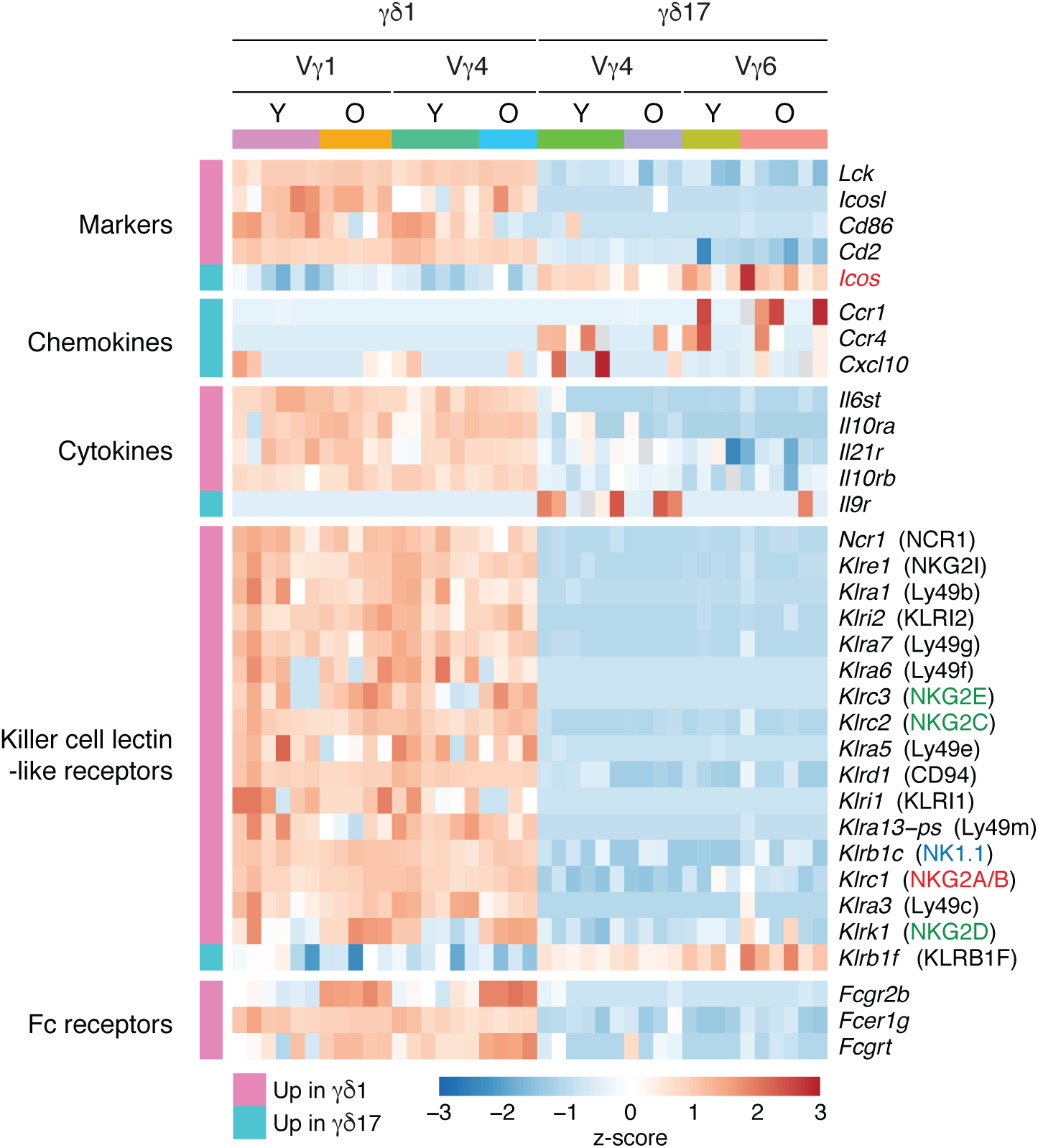
Differentially expressed genes between *γδ*1 and *γδ*17 T cell subsets isolated from young and old mice. Genes expressed at different levels in γδ1 and γδ17 T cells were identified by RNA-Seq analysis. Heatmap shows the Z-score scaled, normalised expression of selected marker genes.

**Suppl. Fig. 6.**
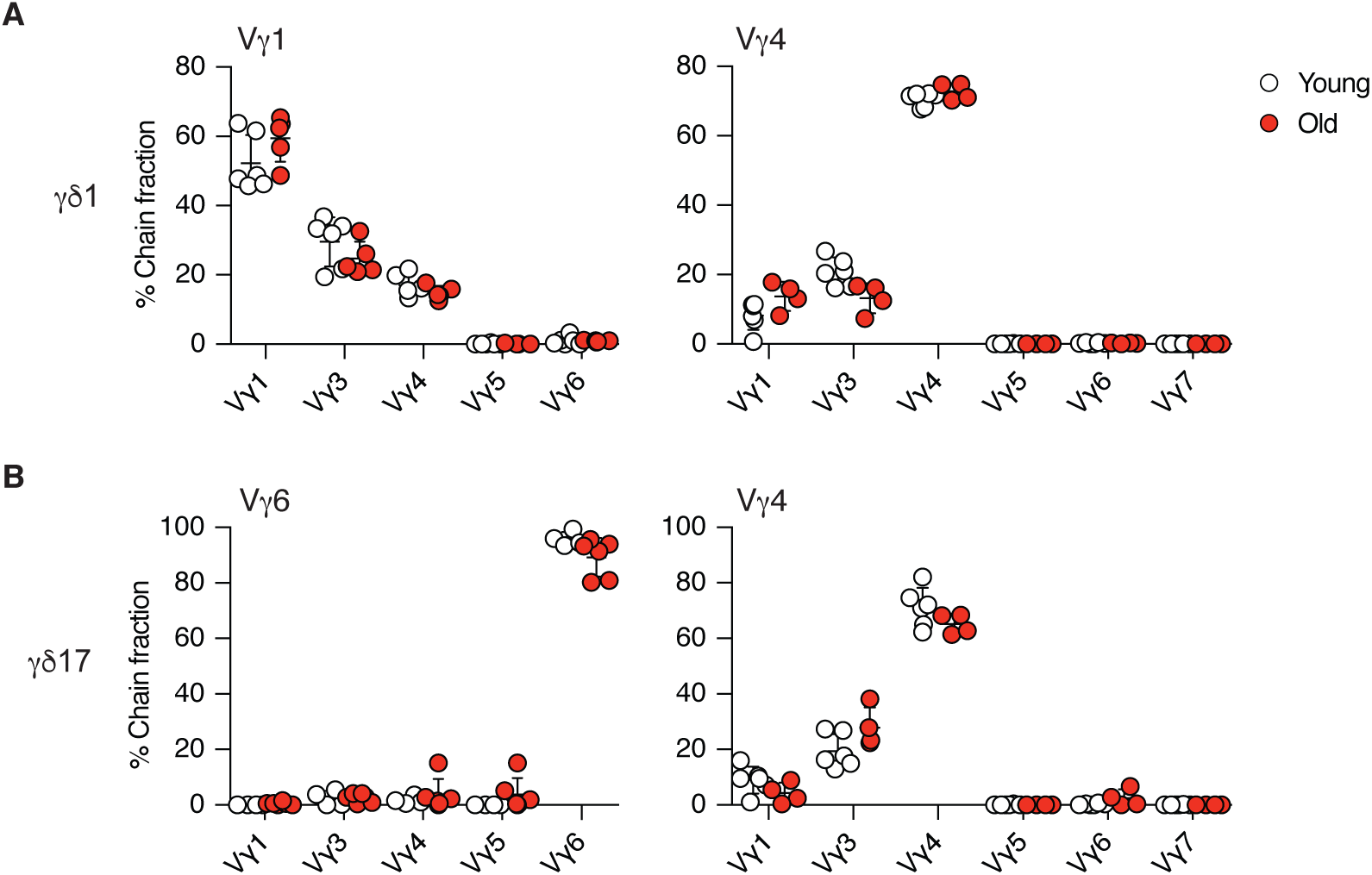
Quality controls of repertoire analysis. TCRγ and TCRδ chains were assembled from bulk RNA-Seq data of highly pure, FACS-sorted **(A)** Vγ1^+^ and Vγ4^+^ γδ1 cells and **(B)** Vγ6^+^ and Vγ4^+^ γδ17 cells. MiXCR was used in RNA-Seq mode to reconstruct TCRγ and TCRδ chains. As a quality control, the fraction of assembled TCRγ chains was plotted for each sample. Only T cell subset-specific sequences were used for further analysis. The detected non-specific sequences are likely due to alignment errors resulting from high level of homology between Vγ1 and Vγ3 (A).

**Suppl. Fig. 7.**
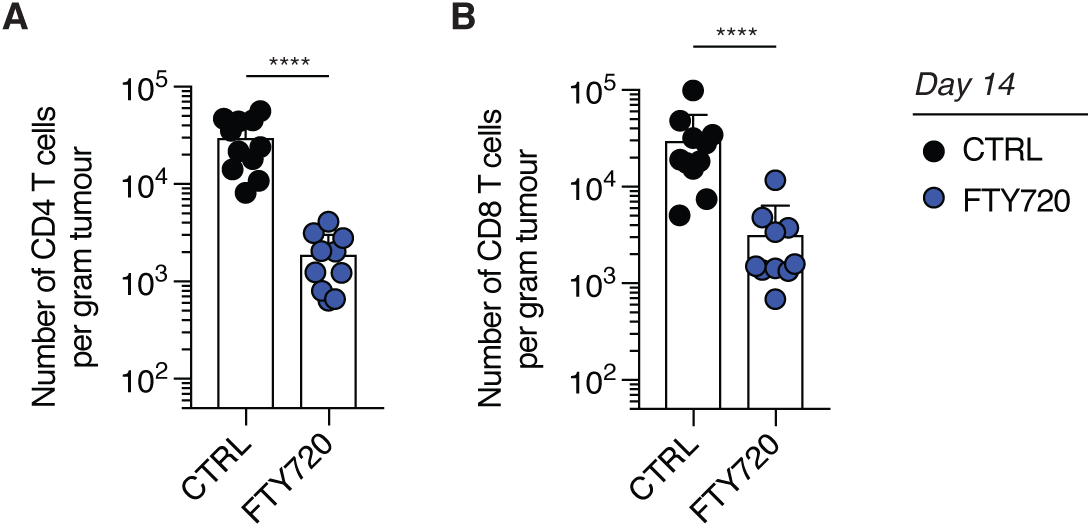
FTY720 treatment blocks egress of *αβ* T cells from the pLNs to the tumour. Young mice were injected every other day by *i.p* with FTY720 (1 mg/kg) or with vehicle control from day -5 to day 13. 3×10^6^ 3LL-A9 cells were injected *s.c* on day 0 and tumours were harvested on day 14 or day 15 for flow cytometry analysis. Number of CD4 **(A)** and CD8 **(B)** T cells in the tumour of control and FTY720-treated mice. Results shown are from 2 independent experiments with 11 control and 10 FTY720-treated mice. Statistical significances for difference in cell densities and cell proportions were assessed by Mann-Whitney test (A and B) or two-way ANOVA (C and D). \*\*\*\**p*<0.0001

**Suppl. Fig. 8.**
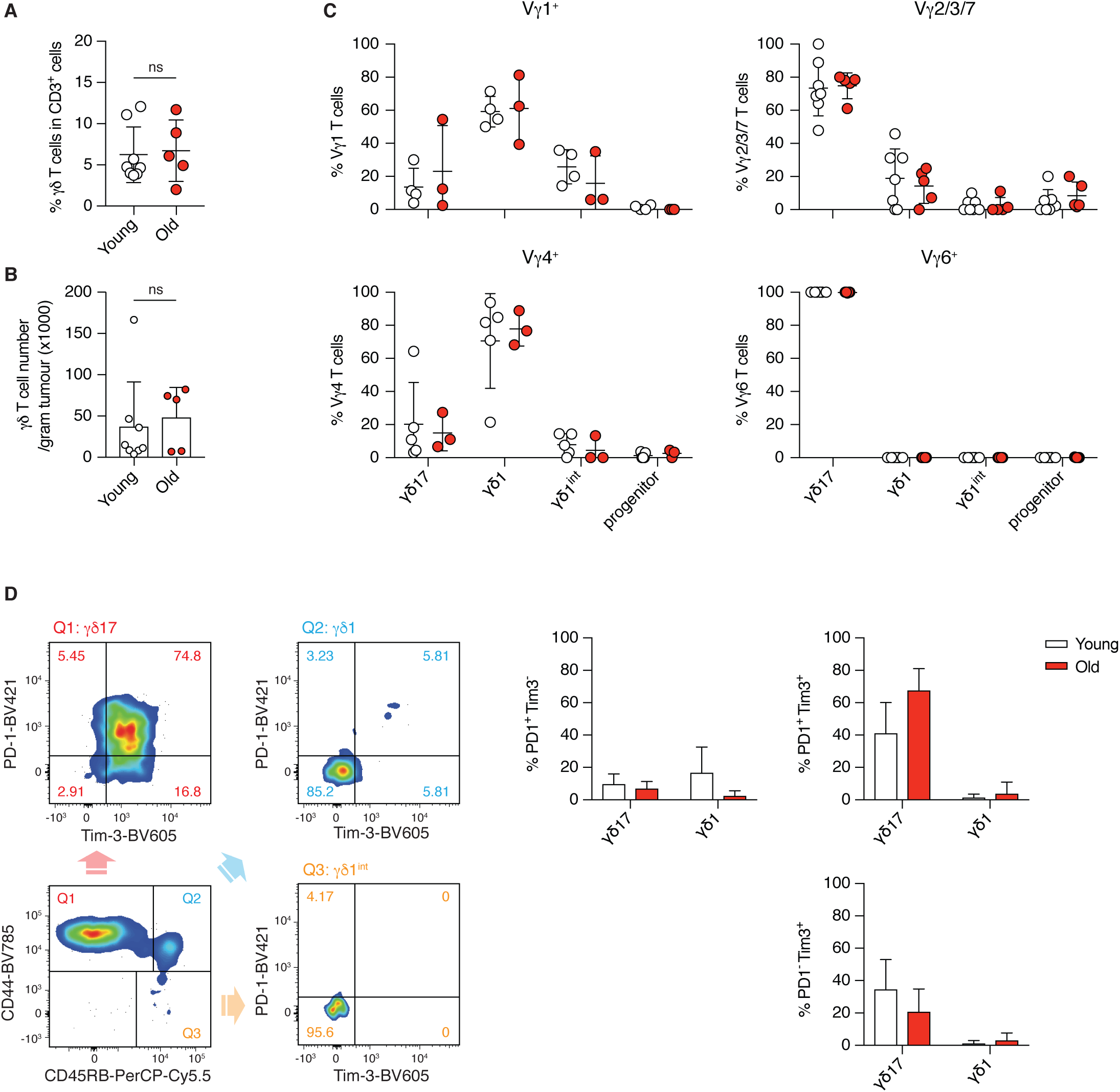
Activation of different *γδ* T cell subsets in the tumour of young and old mice. **(A)** Percentage of γδ T cells in total tumour-infiltrating CD3^+^ T lymphocytes from young and old mice. **(B)** Density of γδ T cells in tumours from young and old mice. **(C)** γδ1 and γδ17 lineage commitment of tumour-infiltrating Vγ1^+^, Vγ2/3/7, Vγ4^+^ and Vγ6^+^ T cells in young and old mice. **(D)** Activation of γδ1 and γδ17 T cell subsets in the tumour is determined by PD-1 and Tim-3 expression. Representative FACS plots (left) show the analysis with concatenated FACS data acquired for each individual young mouse. Results shown (right) are obtained from 2 independent experiments with 8 young and 5 old mice. Cell populations with a total cell number less than 10 were excluded from analysis. Statistical significances for differences were assessed by Mann-Whitney test (A and B) or two-way ANOVA (C and D).

**Suppl. Fig. 9.**
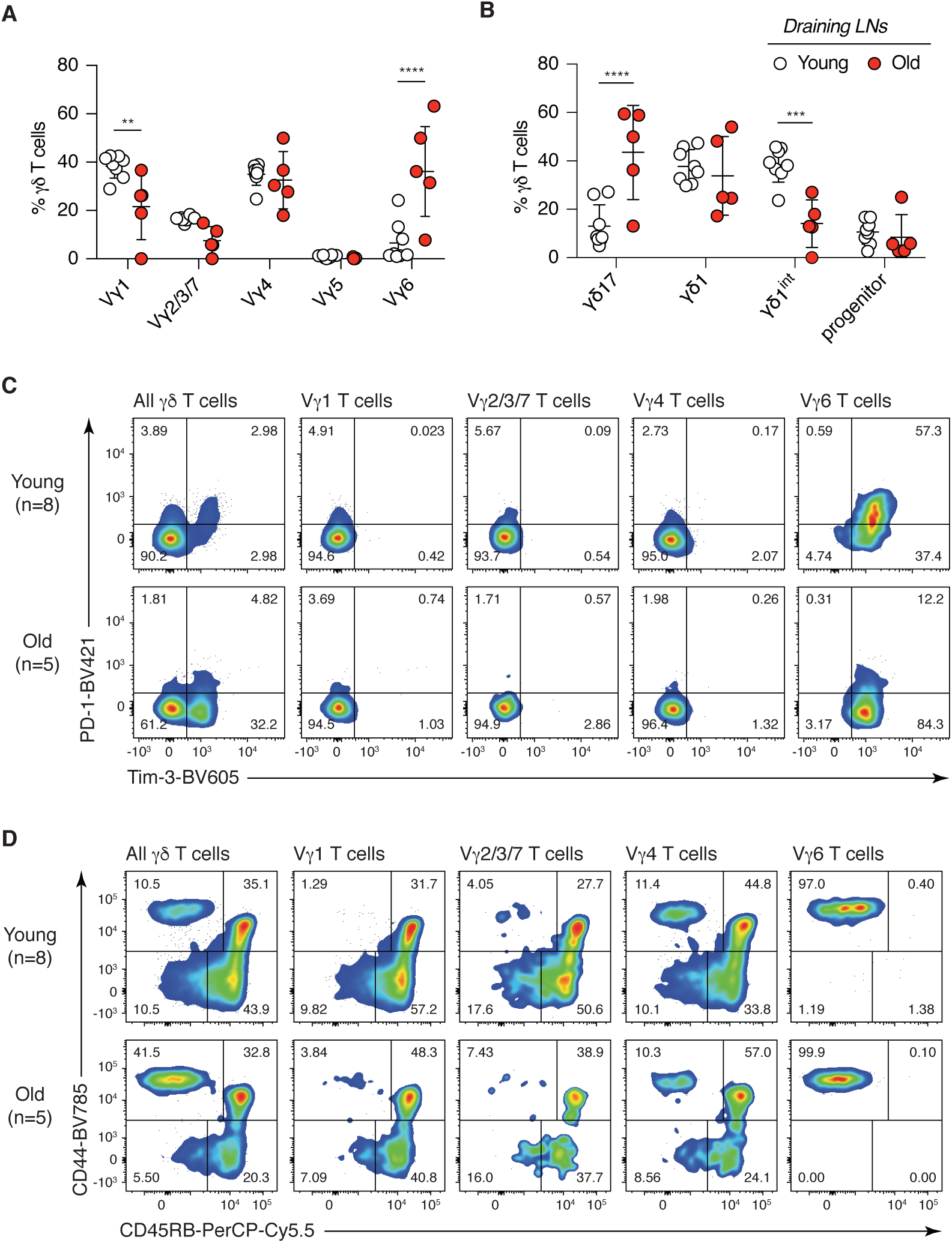
V*γ*6^+^ *γδ*17 T cells are the only *γδ* T cell subset that becomes activated in the draining LN of tumour-bearing young and old mice. **(A)** Proportion of γδ T cell subsets in the tumour-draining LN of young and old mice. **(B)** γδ1 and γδ17 lineage commitment of γδ T cells in the tumour-draining LN of young and old mice. **(C)** Activation status of each γδ T cell subsets in the tumour-draining LN were characterised by PD-1 and Tim-3 expression. **(D)** γδ1 and γδ17 lineage commitment of each γδ T cell subsets in the tumour-draining LN were characterised by PD-1 and Tim-3 expression. FACS files acquired for each individual mouse were concatenated for the analysis and the results are shown as representative dot plots. Results are from 2 independent experiments with 8 young and 5 old mice. Cell populations with the total cell number less than 10 were excluded from analyses. Statistical significances for differences were assessed by two-way ANOVA (A and B). \*\**p*<0.01; \*\*\**p*<0.001; \*\*\*\**p*<0.0001

**Suppl. Fig. 10.**
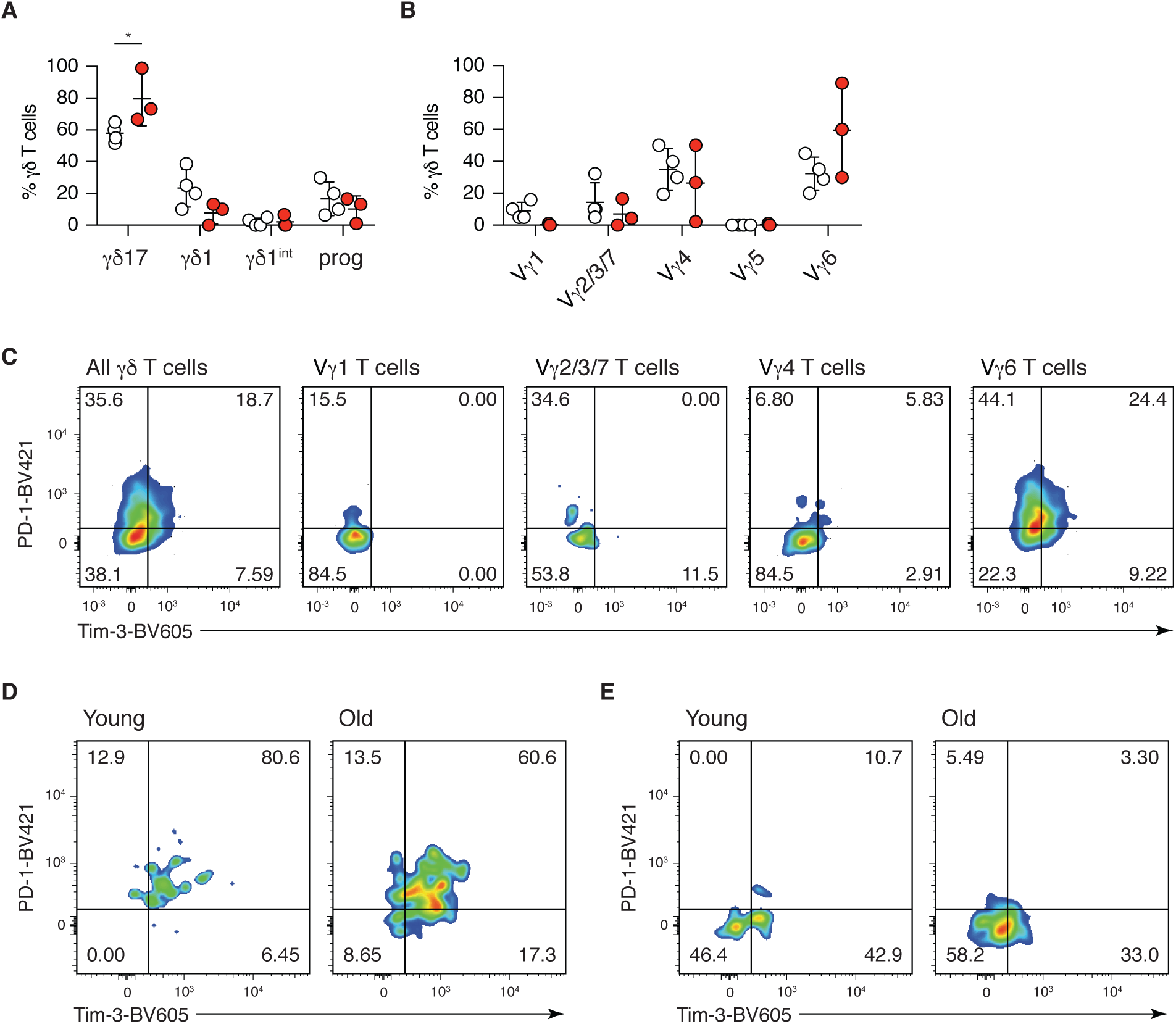
Activation and tumour infiltration of V*γ*6^+^ *γδ*17 T cells in the LL2 tumour model in young and old mice. **(A-E)** Young and old mice were injected subcutaneously with 3×10^6^ LL2 Lewis lung carcinoma cells. Tumour and tumour-draining LN were harvested at indicated times for flow cytometry analysis. **(A)** γδ1 and γδ17 lineage commitment of γδ T cells in the tumour of young and old mice at day 14. **(B)** Proportion of different γδ T cell subsets in the tumour of young and old mice at day 14. **(C)** Activation status of each γδ T cell subsets in the tumour of young mice were characterised by PD-1 and Tim-3 expression at day 11. Activation status of Vγ6^+^ T cells in the tumour **(D)** and tumour-draining LN **(E)** of young and old mice at day 14. FACS files acquired for each individual mouse were concatenated for the analysis and the results were shown as representative dot plots. Results shown in (A, B, D and E) are obtained from an experiment with 4 young and 3 old mice. Results shown in (C) are obtained from an experiments with 5 young mice. Statistical significance for differences was assessed by two-way ANOVA (A and B). \**p*<0.05

**Suppl. Table 1.** Antibodies used in this study

| Antigen | Clone | Dilution | Manufacturer | Identifier | Conjugate |
| --- | --- | --- | --- | --- | --- |
| CD3ε | 145-2C11 | 1:100 | BD | Cat #563565 | BUV395 |
| CD4 | RM4-5 | 1:400 | Biolegend | Cat #100516 | APC |
| CD4 | RM4-5 | 1:50 | Biolegend | Cat #100559 | BV510 |
| CD4 | RM4-5 | 1:200 | Biolegend | Cat #100548 | BV605 |
| CD4 | RM4-5 | 1:200 | Biolegend | Cat #100546 | BV650 |
| CD8α | 53-6.7 | 1:50 | Biolegend | Cat #100752 | BV510 |
| CD8α | 53-6.7 | 1:200 | Biolegend | Cat #100744 | BV605 |
| CD8α | 53-6.7 | 1:200 | Biolegend | Cat #100742 | BV650 |
| CD8α | 53-6.7 | 1:200 | Biolegend | Cat #100748 | BV711 |
| CD8α | 53-6.7 | 1:200 | Biolegend | Cat #100750 | BV785 |
| CD19 | 6D5 | 1:50 | Biolegend | Cat #115546 | BV510 |
| CD24 | M1/69 | 1:200 | Biolegend | Cat #101808 | PE |
| CD24 | M1/69 | 1:200 | Biolegend | Cat #101826 | BV421 |
| CD24 | M1/69 | 1:200 | BD | Cat #563545 | BV650 |
| CD27 | LG.7F9 | 1:100 | eBioscience | Cat #25-0271-82 | PE-Cy7 |
| CD44 | IM7 | 1:200 | Biolegend | Cat #103047 | BV605 |
| CD44 | IM7 | 1:200 | Biolegend | Cat #103059 | BV785 |
| CD45RB | C363-16A | 1:200 | Biolegend | Cat #103314 | PerCP-Cy5.5 |
| CD62L | MEL-14 | 1:400 | Biolegend | Cat #104441 | BV510 |
| CD127 | A7R34 | 1:50 | Biolegend | Cat #135035 | BV711 |
| IL-23R | 12B2B64 | 1:100 | Biolegend | Cat #150904 | PE |
| CCR6 | 140706 | 1:100 | BD | Cat #564736 | BV421 |
| PD-1 | RMP1-30 | 1:50 | Biolegend | Cat #109121 | BV421 |
| PD-1 | RMP1-30 | 1:50 | Biolegend | Cat #109110 | PE-Cy7 |
| Tim-3 | RMT3-23 | 1:50 | Biolegend | Cat #119721 | BV605 |
| IFN-γ | XMG1.2 | 1:50 | Biolegend | Cat #505836 | BV711 |
| IL-17A | TC11-18H10.1 | 1:50 | Biolegend | Cat #506926 | BV421 |
| TCRβ | H57-597 | 1:50 | Biolegend | Cat #109233 | BV510 |
| TCRβ | H57-597 | 1:50 | Biolegend | Cat #109243 | BV711 |
| TCRβ | H57-597 | 1:200 | Biolegend | Cat #109204 | biotin |
| TCR$\gamma\delta$ | UC7-13D5 | 1:100 | Biolegend | Cat #107507 | PE |
| TCRδ | GL3 | 1:100 | Biolegend | Cat #118128 | AF488 |
| TCRδ | GL3 | 1:50 | Biolegend | Cat #118116 | APC |
| TCRδ | GL3 | 1:50 | Biolegend | Cat #118131 | BV510 |
| TCRδ | GL3 | 1:100 | Biolegend | Cat #118129 | BV605 |
| TCR Vγ1 | 2.11 | 1:100 | Biolegend | Cat #141108 | APC |
| TCR Vγ4 | UC3-10A6 | 1:50 | Biolegend | Cat #137704 | FITC |
| TCR Vγ4 | UC3-10A6 | 1:100 | Biolegend | Cat #137704 | PE |
| TCR Vγ5 | 536 | 1:100 | Biolegend | Cat #137506 | APC |
| TCR Vγ5 | 536 | 1:100 | Biolegend | Cat #137504 | PE |
| TCR Vγ5/ Vγ6 | 17D1* | 30 μl/sample | Prof. Adrian Hayday | - | - |
| rat IgM | RM-7B4 | 1:100 | eBioscience | Cat #12-4342-82 | PE |
| CD16/32 (TruStain fcX) | 93 | 1:100 | Biolegend | Cat #101320 | - |

\*hybridoma supernatant

